# Integration of steady-state diffusion MRI with Neural Posterior Estimation (NPE) for post-mortem investigations

**DOI:** 10.1101/2025.07.08.663685

**Authors:** Benjamin C. Tendler

**Affiliations:** Centre for Integrative Neuroimaging, FMRIB, Nuffield Department of Clinical Neurosciences, University of Oxford, Oxford, UK

**Keywords:** Diffusion Weighted Steady-State Free Precession, Neural Posterior Estimation, Simulation Based Inference, Post-Mortem MRI, Diffusion MRI

## Abstract

Post-mortem diffusion MRI plays a key role in investigative pipelines to characterise tissue microstructure, with long scan times facilitating the acquisition of datasets with improved spatial/angular resolution and reduced artefacts versus in vivo. Diffusion-weighted steady-state free precession (DW-SSFP) has emerged as a powerful technique for post-mortem imaging, achieving high SNR-efficiency and strong diffusion weighting in the challenging imaging environment of fixed tissue. However, the sophisticated signal forming mechanisms of DW-SSFP limit the characterisation of tissue properties with conventional parameter estimation routines. Here, I investigate the integration of DW-SSFP with neural posterior estimation (NPE), a parameter inference technique leveraging concepts from Bayesian statistics and machine learning to directly estimate *P*(θ | S) (i.e. the posterior distribution of parameters θ given signal S). A key challenge is that diffusion attenuation in DW-SSFP is dependent on tissue relaxation properties (T_1_/T_2_) and transmit inhomogeneity (B_1_), which must be estimated experimentally and incorporated into the NPE network for accurate modelling. Using a Tensor representation and NPE to estimate *P*(θ | S, *T*_1_, *T*_2_, *B*_1_) (i.e. conditioning on S and known T_1_/T_2_/B_1_), evaluations using synthetic DW-SSP data give excellent agreement to simulation ground truth even in the presence of non-Gaussian noise in low-SNR regimes. Evaluations using experimental DW-SSFP data (whole human post-mortem brain) give excellent agreement to non-linear least-squares estimates, with NPE providing 1000s of posterior samples in a matched evaluation time. Taken together, findings provide a framework to perform rapid parameter estimation with DW-SSFP, and an intuitive approach to incorporate conditional dependencies with NPE.

## 1. Introduction

Non-invasive imaging is a core technology for assessing brain health. Many neuropathologies are characterised by changes at the cellular (μm) scale, motivating the development of imaging techniques sensitive to tissue microstructure. Diffusion MRI is a leading modality in this space, producing images with contrast reflecting the diffusion of water within and surrounding cells (Szafer et al., 1995). Sequence parameters can be adjusted to investigate tissue diffusivity along different spatial orientations (Chenevert et al., 1990; Moseley et al., 1990), length scales, and timing regimes (Bihan, 1995; Vangelderen et al., 1994), with the integration of measured signals with sophisticated biophysical models facilitating the quantitative evaluation of microstructural properties (Alexander et al., 2019; Bihan, 1995; Novikov et al., 2019).

Post-mortem diffusion MRI plays a key role in investigative pipelines to characterise tissue microstructure. Specifically, the long scan times afforded by post-mortem investigations facilitates the acquisition of datasets with improved spatial/angular/b-value resolution and reduced artefacts (e.g. arising from physiological motion) versus in vivo (Roebroeck et al., 2019; Schilling et al., 2025). Post-mortem MRI can also be combined with ultra-high resolution microscopy techniques (performed in the same tissue) to validate microstructural estimates (Choe et al., 2012; Mollink et al., 2017), and directly compared with in vivo MRI to facilitate cross scale-comparisons. More recent work has extended such approaches beyond validation, integrating post-mortem diffusion MRI and microscopy data to address degeneracies (Howard et al., 2019) and improve estimation of brain connectivity (S. Zhu et al., 2025).

A key challenge for post-mortem diffusion MRI investigations is the acquisition of high-quality datasets. Direct translation of in vivo diffusion MRI protocols can lead to images with poor SNR and low diffusion contrast, arising from the short T_2_ and low diffusivity of fixed post-mortem tissue (Roebroeck et al., 2019). Whilst this can be addressed using specialised MRI scanners with powerful gradients, such approaches limit sample size (e.g. pre-clinical) or scanner availability (e.g. CONNECTOM systems) (McNab et al., 2013), motivating investment into acquisition methods more suited to the imaging environment of fixed post-mortem tissue.

Diffusion-weighted steady-state free precession (Kaiser et al., 1974; Le Bihan, 1988; Merboldt et al., 1989b, 1989a) (DW-SSFP) (Figure 1) has emerged as a powerful technique for post-mortem investigations, with signal-forming mechanisms leading to high SNR-efficiency, strong diffusion-weighting, and minimal geometric-distortions (Miller et al., 2012) (see Theory). SNR-efficiency has been demonstrated to increase with magnetic field strength (Foxley et al., 2014), with recent post-mortem DW-SSFP examples including a 500 μm whole human brain (7 T) (Tendler et al., 2022), a 400 μm whole macaque brain (10.5 T human scanner) (Tendler, Warrington, et al., 2025; Warrington et al., 2025), and investigations across a range of human (Cardenas et al., 2017; Edlow et al., 2018; Ji et al., 2024; Nolan et al., 2021; van Veluw et al., 2019; Wilkinson et al., 2016; Q. Zhu et al., 2024) and non-human (Berns et al., 2015; Berns & Ashwell, 2017; Boch et al., 2024; Bryant et al., 2021; Cook & Berns, 2022; Flem et al., 2025; Orekhova et al., 2022; Roumazeilles et al., 2020; Zheng et al., 2024) tissue samples.

**Figure 1:**
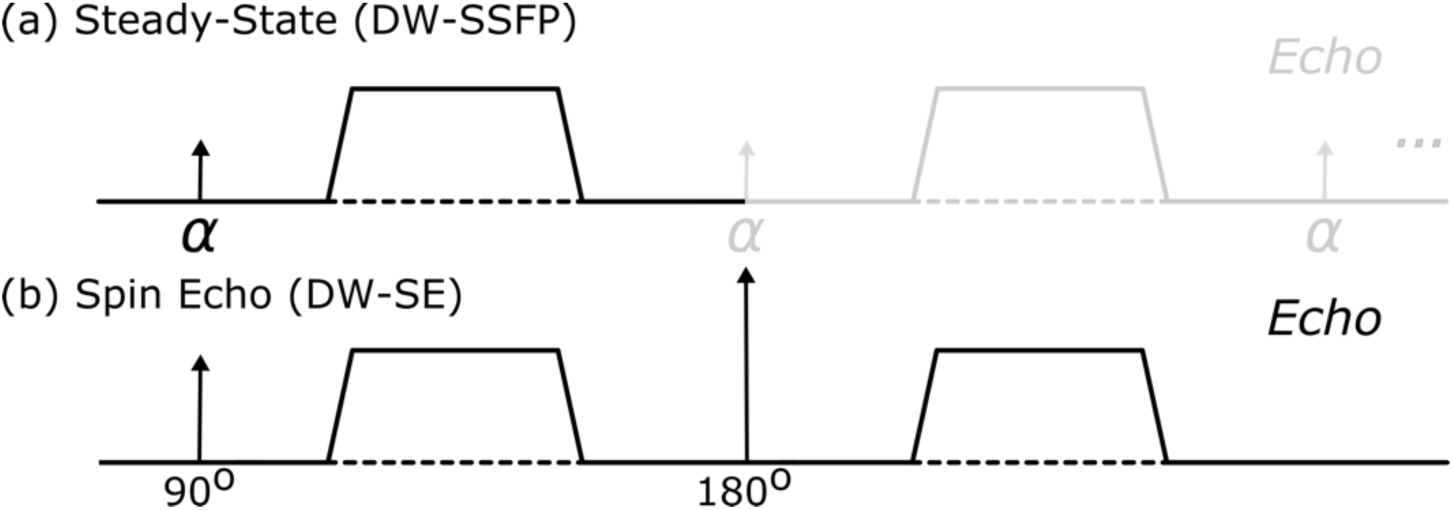
Diffusion encodings. (a) The diffusion encoding of the DW-SSFP sequence consists of a single RF pulse (flip angle *α*) and diffusion gradient per TR (black line). A short TR (∼20 − 40 ms) combined with no spoiling of transverse magnetisation leads to magnetisation being sensitised to diffusion gradients across multiple TRs (grey line). This sensitisation leads to magnetisation dephasing and rephasing consistent with the diffusion encoding of conventional diffusion MRI sequences, e.g. the DW-SE displayed in (b), leading to diffusion-weighted contrast.

DW-SSFP signal measurements have demonstrated distinct microstructural sensitivities versus the diffusion-weighted spin-echo (DW-SE) sequence (McNab & Miller, 2008; Tendler, 2025; Zheng et al., 2024). Whilst previous work has successfully integrated time-independent systems (characterised by Gaussian compartments) with analytical DW-SSFP signal representations for parameter estimation (Foxley et al., 2014; McNab et al., 2009; Tendler, 2025; Tendler, Foxley, Cottaar, et al., 2020), the sophisticated signal-forming mechanisms of DW-SSFP (Freed et al., 2001; Tendler, 2025) currently limits the integration of advanced biophysical models (e.g. incorporating time-dependence or derived from Monte-Carlo simulations) (Novikov et al., 2019) with conventional iterative parameter estimation routines (see Theory). Such extensions would greatly benefit from new data-fitting approaches to support rapid parameter estimation.

In this work, I investigate the integration of DW-SSFP with Neural Posterior Estimation (NPE), a parameter inference technique leveraging concepts from Bayesian statistics and machine learning (Papamakarios & Murray, 2016) to directly estimate the posterior distribution of parameters, *P*(θ | S) (i.e. the probability distribution over parameters θ given data S). Briefly, given a prior distribution of parameters, *P*(θ), and a forward simulation model, *f*(θ), NPE uses simulated data pairs [θ, S] to train a neural network to estimate *P*(θ | S). Once trained, experimental data can be passed to the network to estimate *P*(θ | S_exp_). In comparison to more conventional parameter estimation methods (e.g. non-linear least squares), the estimated posterior distribution can provide detailed information about parameter uncertainties and model degeneracies. A key feature of the implemented NPE network is that inference is *amortised*, i.e. training is performed only once to establish a network capable of performing parameter estimation across voxels in a dataset (see Theory). NPE can be considered a specific case of simulation based inference (SBI), which describes a range of approaches to estimate a target distribution using simulated data (Cranmer et al., 2020).

Previous work has demonstrated the potential of NPE for diffusion-weighted spin-echo (DW-SE) investigations (Eggl & De Santis, 2024; Jallais & Palombo, 2024; Manzano-Patron et al., 2025), providing rapid parameter estimates and detailed posterior distributions to characterise uncertainty and model degeneracies. A key challenge for DW-SSFP is that the signal model has additional voxelwise dependencies on tissue relaxation properties (T_1_/T_2_) and B_1_, which must be estimated experimentally for accurate diffusion modelling (see Theory). From the perspective of NPE, this corresponds to a different relationship between a sample’s diffusivity properties and the measured DW-SSFP signal per T_1_/T_2_/B_1_ combination. Training a different network per T_1_/T_2_/B_1_ combination is infeasible, requiring the adoption of an alternative approach.

Here, I address signal dependences in DW-SSFP by using NPE to estimate *P*(θ | S, *T*_1_, *T*_2_, *B*_1_), i.e. the posterior distribution conditioned on the measured signal and experimentally estimated T_1_, T_2_, and B_1_ values. Using a Tensor representation (Basser et al., 1994) of the DW-SSFP signal, I demonstrate that the proposed network successfully accounts for DW-SSFP signal dependencies, achieving excellent agreement with conventional non-linear least-squares (NLLS) whilst providing 1000s of posterior samples in a matched evaluation time. By additionally incorporating noisy inputs into the training dataset, using synthetic data I demonstrate that the NPE network can achieve excellent agreement to a known ground truth in low SNR regimes characterised by a non-Gaussian noise distribution (i.e. magnitude rectified noncentral chi distribution with a single degree of freedom).

The implemented approach can be adapted to incorporate more sophisticated microstructural models, alternative acquisition schemes, or sequence designs, requiring only a forward simulation model to transform parameters into signal estimates. Experimental evaluations are performed in a DW-SSFP dataset acquired in a whole post-mortem human brain (0.85 mm isotropic resolution). Software, including scripts to replicate many of the findings in this manuscript, is available at github.com/BenjaminTendler/SBI_DWSSFP.

## 2. Theory

### 2.1 Overview of Diffusion-Weighted Steady-State Free Precession (DW-SSFP)

The diffusion encoding of the DW-SSFP sequence (Kaiser et al., 1974; Le Bihan, 1988; Merboldt et al., 1989b, 1989a) consists of a single RF pulse and diffusion gradient per TR (Figure 1a). A short repetition time (TR) (∼20 – 40 ms) combined with no transverse spoiling leads to magnetisation that experiences repeated sensitisation to diffusion gradient pairs over multiple TRs (Figure 1a – grey line), consistent with the dephasing/rephasing of magnetisation associated with the DW-SE sequence (Figure 1b).

From the perspective of extended phase graphs (EPG) (Weigel, 2015), the steady-state DW-SSFP signal can be interpreted as a composite measurement corresponding to the weighted-sum of magnetisation pathways associated with different degrees of diffusion-weighting (Weigel et al., 2010). Whilst this leads to a comparatively complicated set of signal-forming mechanisms versus the DW-SE, these same signal forming mechanisms offer several advantageous properties for diffusion MRI investigations:

- **SNR-efficiency**: A short TR (∼20 – 40 ms) combined with a short diffusion gradient (∼10 ms) leads to a large fraction of each TR dedicated to signal readout and a high SNR-efficiency (Miller et al., 2012).
- **Diffusion-weighting**: Many steady-state magnetisation pathways are associated with long diffusion times or experience repeat sensitisation to diffusion gradients across several TRs, leading to a signal with a high ‘effective’ b-value (Buxton, 1993; Tendler, 2025).
- **Signal sampling**: Sequence timings facilitate a short echo time (TE), advantageous for improving SNR in short T_2_ tissue (Foxley et al., 2014; Miller et al., 2004).
- **Geometric distortion**: The sequence’s short TR is compatible with a single-line readout, achieving low/negligible geometric distortion.

Whilst routine in vivo investigations are currently limited by the sequences severe motion sensitivity (Merboldt et al., 1989b) (an area of continued investigation) (R. O’Halloran et al., 2015; R. L. O’Halloran et al., 2013; Tendler, Wu, et al., 2026), the sequence has become an established method for post-mortem investigations, alongside several ‘low-motion’ in vivo domains (e.g. cartilage, peripheral nerves) (Bieri et al., 2012; Miller et al., 2004; Z. W. Zhang et al., 2008). For an overview of the properties of DW-SSFP, I recommend the review article by McNab & Miller (McNab & Miller, 2010). For a detailed insight into the signal-forming mechanisms of DW-SSFP, I recommend the article by Tendler (Tendler, 2025).

### 2.2 DW-SSFP Signal Dependencies

A key feature of the DW-SSFP sequence is that diffusion-weighting is dependent on several sequence parameters (diffusion gradient amplitude *G*; diffusion gradient duration *δ*, repetition time TR, flip angle *α*), transmit B_1_ (spatially modulating the flip angle *α*), and sample properties complementary to diffusivity (T_1_ and T_2_). From the perspective of EPG, these dependencies can be interpreted as modifying the relative weighting of magnetisation pathways with different degrees of diffusion weighting (Weigel et al., 2010).

Figure 2 motivates this concept visually, displaying (a) two DW-SSFP signals with identical profiles, but distinct T_1_, T_2_, B_1_, and diffusivity properties and (b) the dependency of diffusion attenuation on T_1_, T_2_ and B_1_, where I note that the dependency relationship will be modulated by the choice of DW-SSFP sequence parameters. To estimate diffusion coefficients, these dependencies must be precisely accounted for when performing signal modelling. For dependencies that vary spatially across a dataset (T_1_, T_2_ and B_1_), this requires the acquisition of complementary experimental MRI data using quantitative T_1_, T_2_ and B_1_ mapping techniques. Given the lengthy scan times associated with post-mortem diffusion MRI protocols, acquisition of these complementary maps typically takes a fraction of the total experimental time, and is commonplace in DW-SSFP investigations (McNab et al., 2009; Miller et al., 2012; Tendler, Foxley, Hernandez-Fernandez, et al., 2020).

**Figure 2:**
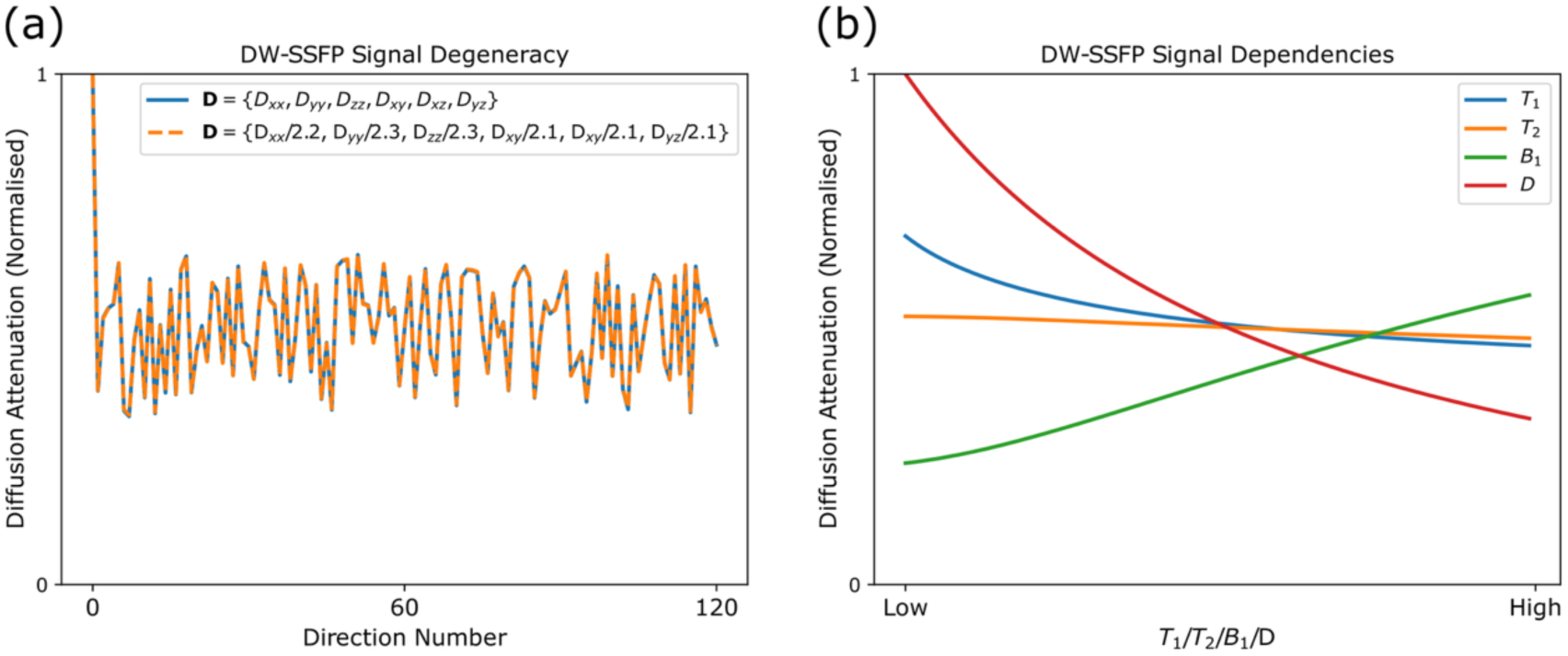
DW-SSFP signal degeneracy and dependencies. (a) Using a Tensor representation of diffusivities, here I display two simulated DW-SSFP signals with distinct diffusion coefficients but identical diffusion attenuation profiles. Profile degeneracy arises from the dependencies of the DW-SSFP signal on T_1_, T_2_ and B_1_, which additionally modulates the diffusion-weighting of DW-SSFP. Specifically, the two DW-SSFP signals have distinct T_1_ (blue: 650 ms; orange: 800 ms), T_2_ (blue: 35 ms; orange: 60 ms) and B_1_ (blue: 1; orange: 0.5) values, which in combination with differing diffusion coefficients leads to an identical signal profile. (b) Dependency of the DW-SSFP signal on T_1_, T_2_ and B_1_ (blue, orange and green lines) modulates diffusion attenuation similar to a change in the diffusion coefficient (red), with the relationship dependent on the choice of DW-SSFP sequence parameters. Taken together, incorporation of T_1_, T_2_ and B_1_ estimates (using complementary mapping techniques) is required for precise estimation of diffusion coefficients in DW-SSFP with a conventional acquisition scheme. Simulations performed using the experimental acquisition parameters described in the Methods at a single flip angle (*α* = 24°). Default diffusion coefficients in (a) based on Tensor estimates in a single corpus callosum voxel of experimental post-mortem data, defining ***D*** = {1.63, 0.81, 0.82, −0.41, −0.40, 0.22} ⋅ 10^−4^ mm^2^/s. Default values in (b) correspond to the mean relaxation values estimated in the acquired post-mortem experimental data and no transmit inhomogeneity (T_1_ = 650 ms; T_2_ = 35 ms; B_1_ = 1), fixing *D* = 1 ⋅ 10^−4^ mm^2^/s.

### 2.3 DW-SSFP Signal Modelling

The complexity of the signal-forming mechanisms of DW-SSFP is reflected in Appendix 1, which presents an analytical expression for the DW-SSFP signal incorporating free Gaussian diffusion (Freed et al., 2001). In contrast to a simple mono-exponential (*S*_0_ ⋅ *e*^−*bD*^) signal expression for the DW-SE sequence, the DW-SSFP sequence is described using a detailed recursive expression.

Whilst more computationally demanding, the expression in Appendix 1 is not a barrier itself to the integration of parameter estimation routines with DW-SSFP data. Previous studies have successfully incorporated a (1)

Tensor (Foxley et al., 2014), (2) Ball & Sticks (McNab et al., 2009), and (3) Gamma distribution of diffusivities (Tendler, Foxley, Cottaar, et al., 2020) to characterise tissue properties. More recent work introduced an alternative representation of the DW-SSFP signal to facilitate the incorporation of biophysical models (Tendler, 2025), establishing a data fitting approach to obtain NODDI (H. Zhang et al., 2012) estimates from experimental DW-SSFP data. To date, parameter estimation has been perform using iterative NLLS or Markov Chain Monte Carlo (MCMC) methods.

When considering biophysical models with time-dependent features (e.g. arising from restrictions experienced by mobile spins), forward-simulations can be performed using (1) Monte Carlo simulations or (2) the Gaussian Phase Approximation (GPA) (Stepišnik, 1993; Tendler, 2025). The long times required to perform a single forward simulation using these methods for DW-SSFP limits their integration with iterative parameter estimation routines. Specifically, Monte Carlo simulations require the DW-SSFP signal to reach a steady state over many TRs, increasing simulation time versus an equivalent DW-SE investigation, as described in (Tendler, 2025). GPA estimations must be averaged over many signal-forming pathways, requiring extensive parallelisation or limiting parameter estimation to approximation regimes of the DW-SSFP signal (Tendler, 2025). The sequence’s dependencies on T_1_, T_2_ and B_1_ can necessitate extremely large storage and simulation requirements using conventional dictionary-based fitting methods.

I propose to address this by integrating DW-SSFP with NPE for rapid parameter inference, as described in the next section.

### 2.4 Neural Posterior Estimation (NPE)

NPE leverages advances in machine learning network architectures to estimate *P*(θ | S) (i.e. the probability distribution over parameters θ given data S). In contrast to parameter estimation procedures that provide pointwise estimates of θ (e.g. non-linear least squares) and uncertainty (e.g. Hessian-based approximations), NPE supports the characterisation of parameter estimates, uncertainty, and degeneracies using detailed posterior distributions (Cranmer et al., 2020; Jallais & Palombo, 2024). Implemented NPE networks facilitate the rapid and direct characterisation of multidimensional posterior distributions with high sampling density, in contrast to conventional posterior-estimation procedures typically associated with reduced sampling efficiency (e.g. approximate Bayesian computation) or iterative sampling schemes (e.g. MCMC) (Cranmer et al., 2020).

NPE network training is performed using simulated data, synthesising many parameter/data pairs ([θ, S]) estimated from a prior distribution, *P*(θ), and a forward simulation model, *f*(θ). Once trained, experimental data (S_exp_) can be passed to the network to directly estimate *P*(θ | S_exp_). Whilst the training phase is typically slow (e.g. several hours/days), inference can be amortised (i.e. training only needs to be performed once) and sampling the posterior distribution using the trained network is extremely fast (e.g. milliseconds per voxel).

In this work, I apply NPE using an invertible neural network architecture incorporating normalising flows (Papamakarios et al., 2021) via a neural spline flows (NSF) density estimator (Durkan et al., 2019). Specifically, the network estimates the transformation between a base (e.g. a standard normal distribution, as used in this manuscript (Durkan et al., 2019)) PDF and *P*(θ | S), with Figure 3a providing a high-level overview of the network design. This approach aims to preserve *P*(θ | S) as a valid PDF by maintaining key properties of the base PDF as it is transformed through the network (e.g. preserving volume changes), whilst providing a sophisticated transformation approach capable of approximating detailed posterior distributions (Durkan et al., 2019; Papamakarios et al., 2021).

**Figure 3:**
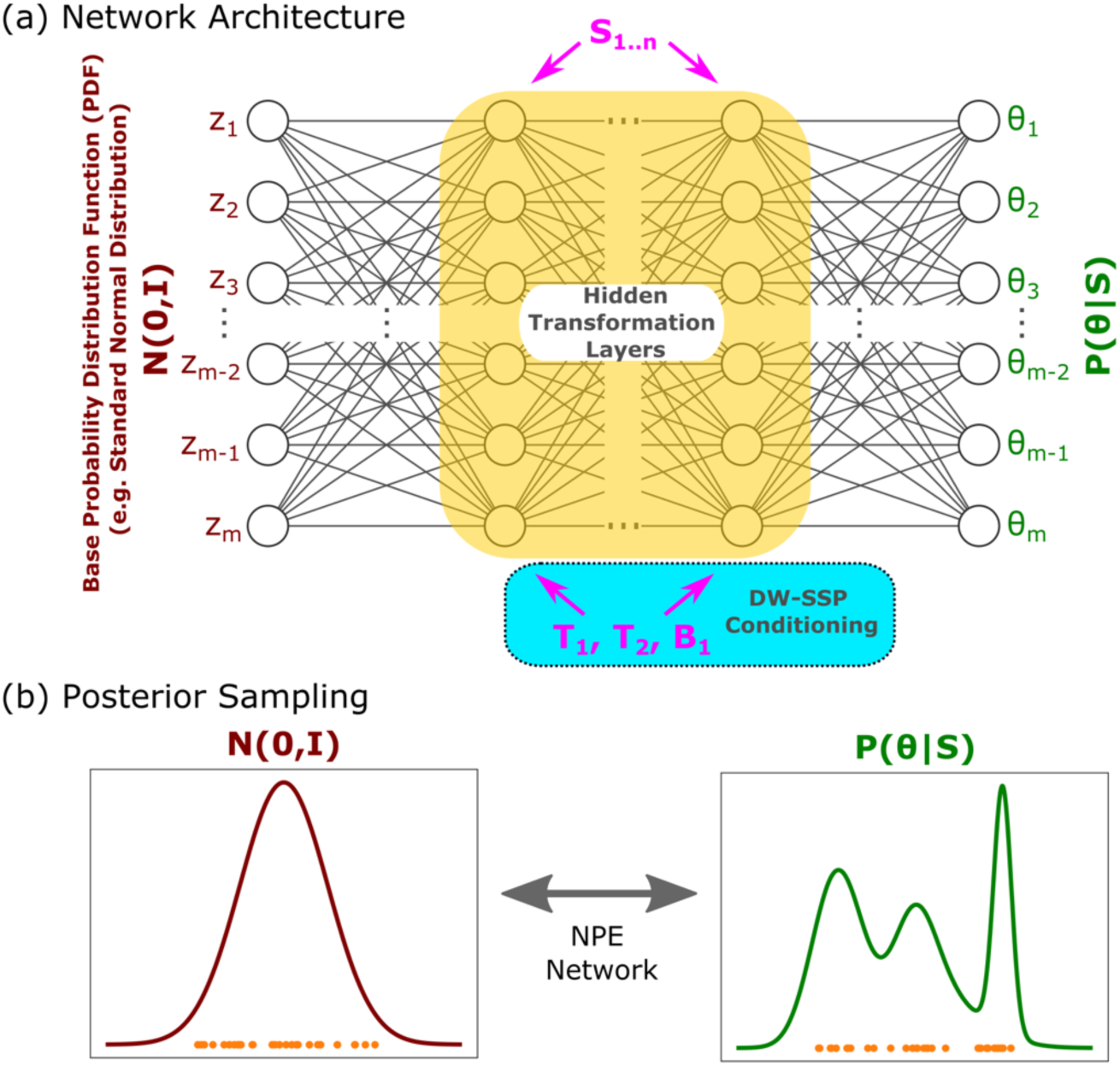
NPE network architecture incorporating DW-SSFP signal dependencies. (a) High-level overview of the network architecture in NPE implemented with normalising flows (Papamakarios et al., 2021), an invertible network learning the transformation between a base PDF (left), and *P*(θ | S) (right). Conditioning is achieved by passing S (dimensions 1 x n) to the ‘hidden’ transformation network layers (yellow), where I propose additionally passing T_1_, T_2_, and B_1_ for DW-SSFP conditioning (blue). (b) Once trained, posterior samples, *θ*∼*P*(θ | S), are generated by transforming base samples, *Z*∼*N*(0, *I*), through the network. Samples of Z and θ visualised in (b) as orange dots, with the density of sampled points reflecting the PDF.

Once trained, samples from the posterior, *θ*∼*P*(θ | S), can be generated by sampling from the base PDF, *Z*∼*N*(0, *I*) and passing the samples through the network (Figure 3b). Conditioning is achieved by additionally passing S to the network (i.e. via ‘hidden’ transformation network layers) (Figure 3a). NPE methods have been shown to precisely approximate sophisticated PDFs (Durkan et al., 2019), with previous work with the DW-SE sequence demonstrating accurate parameter inference for a range of microstructural models (Eggl & De Santis, 2024; Jallais & Palombo, 2024; Manzano-Patron et al., 2025). For further details on normalising flows and NPE, I recommend the articles by Papamakarios et al. (Papamakarios et al., 2021), and Radev et al. (Radev et al., 2023).

To account for dependencies of the DW-SSFP signal on T_1_, T_2_ and B_1_ (see above Theory), I propose additionally passing experimental estimates of T_1_, T_2_ and B_1_ to the hidden transformation network layers for conditioning (Figure 3a - blue box). This approach approximates the posterior PDF as *P*(θ | S, *T*_1_, *T*_2_, *B*_1_), avoiding the requirement to train a separate network per instance of T_1_, T_2_ and B_1_.

## 3. Methods

### 3.1 DW-SSFP forward model

A forward model of the DW-SSFP signal under a Tensor representation was established incorporating DW-SSFP acquisition parameters, T_1_, T_2_, B_1_, and Tensor coefficients (Appendix 2), implemented in Python (3.12.8). DW-SSFP sequence parameters were defined based on the experimental DW-SSFP dataset analysed in this manuscript, provided in Table 1. For full details motivating the choice of sequence parameters, see (Tendler, Foxley, Hernandez-Fernandez, et al., 2020).

**Table 1:**
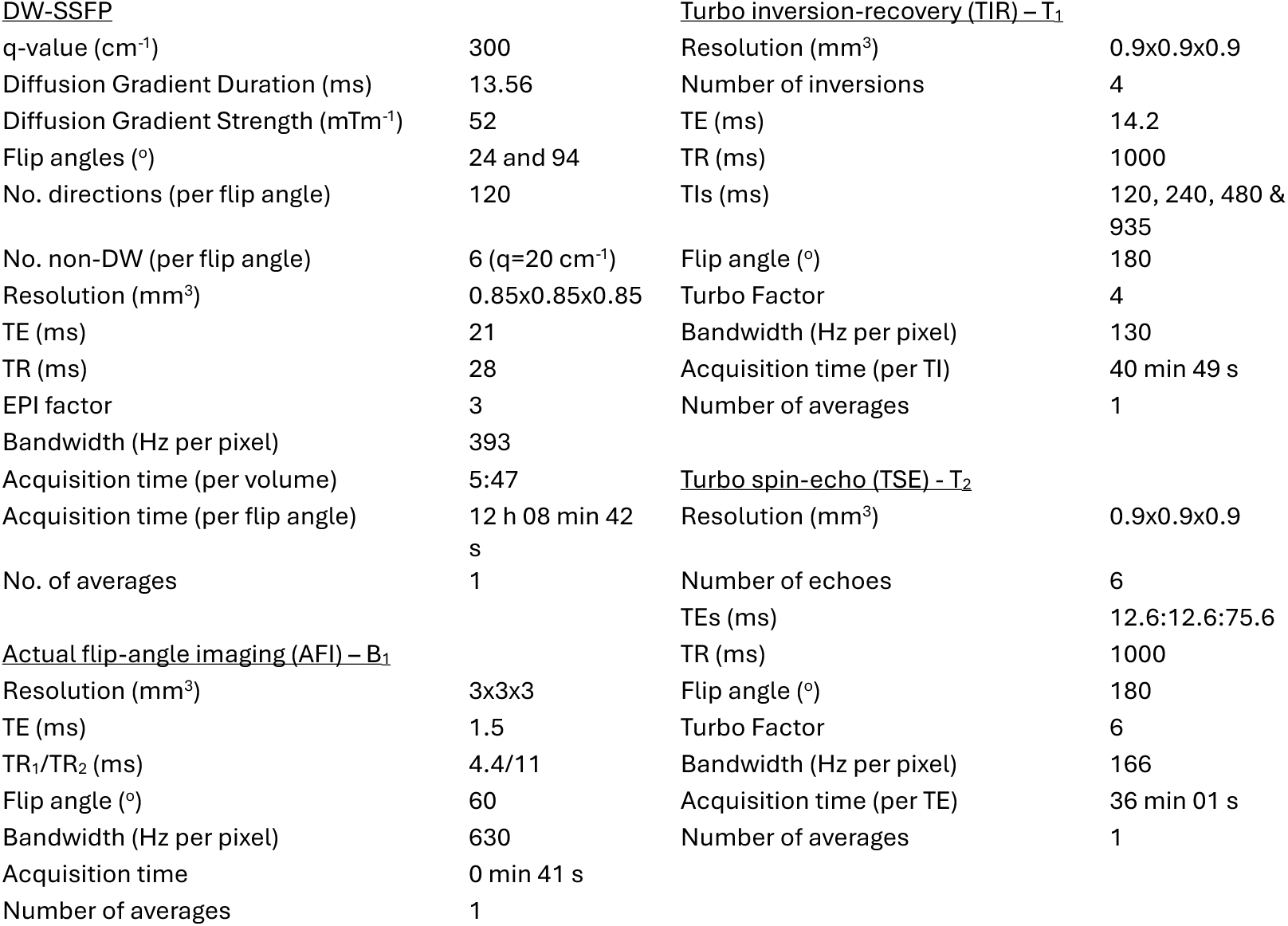
Experimental acquisition parameters. Acquisition parameters for the DW-SSFP sequence and complementary data acquired for quantitative T_1_, T_2_ and B_1_ mapping. For full details motivating the choice of sequence parameters, see (Tendler, Foxley, Hernandez-Fernandez, et al., 2020).

|  |  |  |  |
| --- | --- | --- | --- |
| <u>DW-SSFP</u> |  | <u>Turbo inversion-recovery (TIR) – $T_1$</u> |  |
| q-value ($\text{cm}^{-1}$) | 300 | Resolution ($\text{mm}^3$) | 0.9x0.9x0.9 |
| Diffusion Gradient Duration (ms) | 13.56 | Number of inversions | 4 |
| Diffusion Gradient Strength ($\text{mTm}^{-1}$) | 52 | TE (ms) | 14.2 |
| Flip angles ($^\circ$) | 24 and 94 | TR (ms) | 1000 |
| No. directions (per flip angle) | 120 | TIs (ms) | 120, 240, 480 & 935 |
| No. non-DW (per flip angle) | 6 ($q=20 \text{ cm}^{-1}$) | Flip angle ($^\circ$) | 180 |
| Resolution ($\text{mm}^3$) | 0.85x0.85x0.85 | Turbo Factor | 4 |
| TE (ms) | 21 | Bandwidth (Hz per pixel) | 130 |
| TR (ms) | 28 | Acquisition time (per TI) | 40 min 49 s |
| EPI factor | 3 | Number of averages | 1 |
| Bandwidth (Hz per pixel) | 393 |  |  |
| Acquisition time (per volume) | 5:47 | <u>Turbo spin-echo (TSE) – $T_2$</u> |  |
| Acquisition time (per flip angle) | 12 h 08 min 42 s | Resolution ($\text{mm}^3$) | 0.9x0.9x0.9 |
| No. of averages | 1 | Number of echoes | 6 |
|  |  | TEs (ms) | 12.6:12.6:75.6 |
| <u>Actual flip-angle imaging (AFI) – $B_1$</u> |  | TR (ms) | 1000 |
| Resolution ($\text{mm}^3$) | 3x3x3 | Flip angle ($^\circ$) | 180 |
| TE (ms) | 1.5 | Turbo Factor | 6 |
| TR<sub>1</sub>/TR<sub>2</sub> (ms) | 4.4/11 | Bandwidth (Hz per pixel) | 166 |
| Flip angle ($^\circ$) | 60 | Acquisition time (per TE) | 36 min 01 s |
| Bandwidth (Hz per pixel) | 630 | Number of averages | 1 |
| Acquisition time | 0 min 41 s |  |  |
| Number of averages | 1 |  |  |

### 3.2 Network Implementation

All data analysis was performed in Python (3.12.8), with the conditioned NPE network implemented using the SBI toolbox (0.23.3) (Boelts et al., 2024; Tejero-Cantero et al., 2020) as follows:

- *Priors*: Tensor coefficients (**D**), uniform distributions with limits:

∘ [0, 5] ⋅ 10^−4^ mm^2^/s for diagonal components (*D_xx_*, *D_yy_*, *D_zz_*)
∘ [-2.5, 2.5] ⋅ 10^−4^ mm^2^/s for off-diagonal components (*D_xy_*, *D_xz_*, *D_yz_*)
- *Data Pairs*: 6,000,000 data pairs, corresponding to 1,000,000 simulations with 5 noise levels + noise free. Simulated signals corresponded to a 1D array of dimensions 1×252 (Table 1), with each simulation associated with an arbitrary:

∘ Tensor (sampled from the prior)
∘ T_1_ (uniform distribution, limits = [300, 1200] ms)
∘ T_2_ (uniform distribution, limits = [20, 80] ms)
∘ B_1_ (uniform distribution, limits = [0.2, 1.2])
∘ SNR (uniform distribution, limits = [2,50]). SNR was defined with respect to the non-diffusion weighted (i.e. q = 20 cm^-1^) signal amplitude, setting 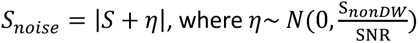. This reflects a magnitude rectified noncentral chi distribution with a single degree of freedom, used to evaluate the capacity of NPE to address parameter biases commonly associated with non-Gaussian noise.
- *Training*: The NPE network was implemented using an NSF density estimator (Durkan et al., 2019), with default SBI toolbox parameters: NPE-C/APT algorithm (Greenberg et al., 2019); no. hidden features = 50, no. transforms = 5; no. spline bins = 10; parameter and data z-score: independent; one training round. T_1_, T_2_ and B_1_ conditioning was achieved by appending their values to the signal during training, creating a 1D array corresponding to [S, T_1_, T_2_, B_1_] of dimensions 1×255.

Similar to the approach in (Manzano-Patron et al., 2025), an additional classifier network was trained to identify regions of the prior distribution (i.e. parameter combinations) that produced positive semi-definite tensors, implemented using the *RestrictionEstimator* function in the SBI toolbox (ResNet; no. hidden features = 100, no. blocks = 2; dropout prob. = 0.5; parameter z-score: independent) (Deistler et al., 2022; Lueckmann et al., 2017). This can be interpreted as modifying the prior distribution prior to synthesising data pairs to avoid invalid parameter combinations. The classifier network was trained using 500,000 simulated data pairs, with the resulting prior distribution visualised in Supplementary Material Figure S1. During training of the classifier network, 53.7% of the 500,000 data pairs were rejected, with incorporation of the trained classifier network accelerating data synthesis by 134.7%. Following training of the classifier network, ∼7.3% of the 6,000,000 data pairs synthesised for NPE network training were rejected.

The total training time was ∼34 hours on a personal laptop (Macbook Pro, Sonoma 14.5, M1, 16GB RAM) with a CPU. This consisted of ∼33 hours for the NPE network training and < 1 hour to train the classifier network and synthesise the noisy data pairs. The trained NPE network required 1.1 MB in storage space.

### 3.3 Network Evaluation: Synthetic data

To evaluate the performance of the conditioned NPE network, simulated DW-SSFP signals were passed to the trained network for parameter inference. **D** was subsequently estimated from the simulated signals and compared to the known ground truth. Two investigations were performed to estimate parameters for (1) fixed **D** across a range of T_1_, T_2_ and B_1_ values, and (2) arbitrary **D**, T_1_, T_2_ and B_1_ combinations. 100 posterior samples were taken per simulation, with the mean of the posterior estimates used for evaluation.

To evaluate the effectiveness of the proposed conditioning strategy for precisely estimating **D** for signals with different relaxation times and transmit inhomogeneity profiles, I performed comparisons with an identical network trained with DW-SSFP signals associated with a single relaxation time (T_1_ = 650 ms and T_2_ = 35 ms, the mean of the experimental post-mortem dataset) and no transmit inhomogeneity (B_1_ = 1).

To characterise the network’s robustness to data noise and parameter estimation under a non-Gaussian noise distribution, a similar evaluation was performed as a function of SNR. **D** was estimated and compared to the known ground truth from simulated DW-SSFP signals with added non-Gaussian noise for the NPE network as described above and an identical NPE network trained with simulated data incorporating a Gaussian noise distribution. Two investigations were performed to estimate parameters for (1) fixed **D** across a range of SNR levels (5 to 50), and (2) arbitrary **D**, T_1_, T_2_ and B_1_ combinations in a low SNR regime (SNR = 5). 100 posterior samples were averaged per NPE simulation, with parameter estimates additionally averaged over 1000 noise realisations per SNR level (NPE and NLLS) for the fixed **D** investigation.

To assess the speed of NPE for parameter estimation, comparisons were made to NLLS as a function of the number of posterior samples, evaluating 1000 arbitrary **D**, T_1_, T_2_ and B_1_ combinations with SNR = 20. Here, the NLLS fitting implementation was performed SciPy *curve_fit* (1.12.0) (trust region reflective algorithm) (E. Jones et al., 2001). Finally, the number of samples required to robustly estimate the mean and standard deviation of the Tensor coefficients for the implemented DW-SSFP acquisition scheme (Table 1) was assessed as a function of SNR for 1000 arbitrary **D**, T_1_, T_2_ and B_1_ combinations.

### 3.4 Network Evaluation: Experimental data

Experimental DW-SSFP and T_1_/T_2_/B_1_ mapping data acquired in a single whole human post-mortem brain on a 7T Siemens system (32ch-receive/1ch-transmit head coil) were used for investigation. These data were acquired as part of an existing project investigating the pathology of amyotrophic lateral sclerosis (ALS) in the human brain (Pallebage-Gamarallage et al., 2018). Motivation behind the choice of acquisition protocol (Table 1) is described in detail in (Tendler, Foxley, Hernandez-Fernandez, et al., 2020), with dependency maps visualised in Supplementary Material Figure S2. Post-mortem tissue was provided by the Oxford Brain Bank, a research ethics committee (REC) approved, human tissue act (HTA) regulated research tissue bank. Research was conducted under the Oxford Brain Bank’s generic Research Ethics Committee approval (15/SC/0639).

Prior to fitting, experimental data were pre-processed following the procedure in (Tendler, Foxley, Hernandez-Fernandez, et al., 2020). Briefly, a quantitative T_1_ map was estimated from the TIR data assuming mono-exponential signal recovery. A quantitative T_2_ map was estimated from the TSE data using an EPG signal model as previously described (Tendler et al., 2021). A quantitative B_1_ map was estimated from the AFI data following the processing steps in the original manuscript (Yarnykh, 2007). The T_1_, T_2_ and B_1_ maps were coregistered to the DW-SSFP data using a 6 degrees of freedom (DOF) transformation with FSL FLIRT (Jenkinson et al., 2002; Jenkinson & Smith, 2001). Whilst the DW-SSFP data was acquired with an EPI factor of 3 (three lines of k-space were acquired per TR), this low EPI factor leads to DW-SSFP volumes with negligible image distortions, meaning no correction for geometric distortion is required. This is motivated in Supplementary Material Figure S2, which visualises the DW-SSFP data alongside quantitative T_1_, T_2_ and B_1_ maps, where alignment has been performed using a six degrees of freedom coregistration only.

For the DW-SSFP data, a Gibbs ringing correction was first applied (Kellner et al., 2016). Individual volumes were subsequently coregistered using a 6 DOF transformation with FSL FLIRT. The DW-SSFP magnitude data was reconstructed using a root sum-of-squares coil combination, leading to a noncentral chi noise distribution (Aja-Fernández et al., 2009). To reduce parameter bias in low SNR regions (D. K. Jones & Basser, 2004) and facilitate comparisons between NPE and NLLS, a noise floor correction was performed to the experimental data (Gudbjartsson & Patz, 1995), with the noise floor estimated from the background signal. To circumvent the requirement of estimating the signal amplitude (S_0_) as part of the parameter estimation procedure, S_0_ was estimated from the non-diffusion weighted (q = 20 cm^-1^) volumes using the expression in Appendix 1 and removed from the experimental data (via division) prior to fitting.

The DW-SSFP signal, alongside T_1_, T_2_ and B_1_ maps were passed to the NPE network on a voxelwise basis to estimate *P*(**D** | S, *T*_1_, *T*_2_, *B*_1_), with 100 posterior samples estimated per voxel. The mean of the posterior samples were used for comparison to the NLLS fitting implementation. In a small fraction of voxels (∼1.3% of the dataset), Tensor coefficients fell outside of the prior bounds (> 5 ⋅ 10^−4^ mm^2^/s) and could not be estimated using NPE. This predominantly corresponded to voxels at the brain surface associated with residual fixative trapped within brain sulci or ventricles, and were not incorporated into the remaining analysis.

## 4. Results

### 4.1 Simulations

Figure 4a-c compares estimated Tensor coefficients as a function of (a) T_1_, (b) T_2_, and (c) B_1_ using NPE for fixed **D** with an NPE network trained with the proposed conditioning approach (left) in comparison to an NPE network trained with fixed relaxation times and no transmit inhomogeneity (right). When incorporating the proposed conditioning approach (right), excellent agreement is found versus ground truth (median absolute difference = 5.0 ·10^-7^ mm^2^/s; interquartile range = 8.1 ·10^-7^ mm^2^/s across the range of T_1_, T_2_, and B_1_, demonstrating that the implemented NPE network can account for DW-SSFP signal dependencies. For the NPE network trained with fixed relaxation times and no transmit inhomogeneity (right), considerable deviations are observed from ground truth, most notably for dependencies on T_1_ and B_1_, consistent with the attenuation trends in Figure 2b. To aid visualisation, Supplementary Material Figure S3 displays the difference between estimated Tensor coefficients in comparison to the simulated ground truth for Figure 4a-c.

**Figure 4:**
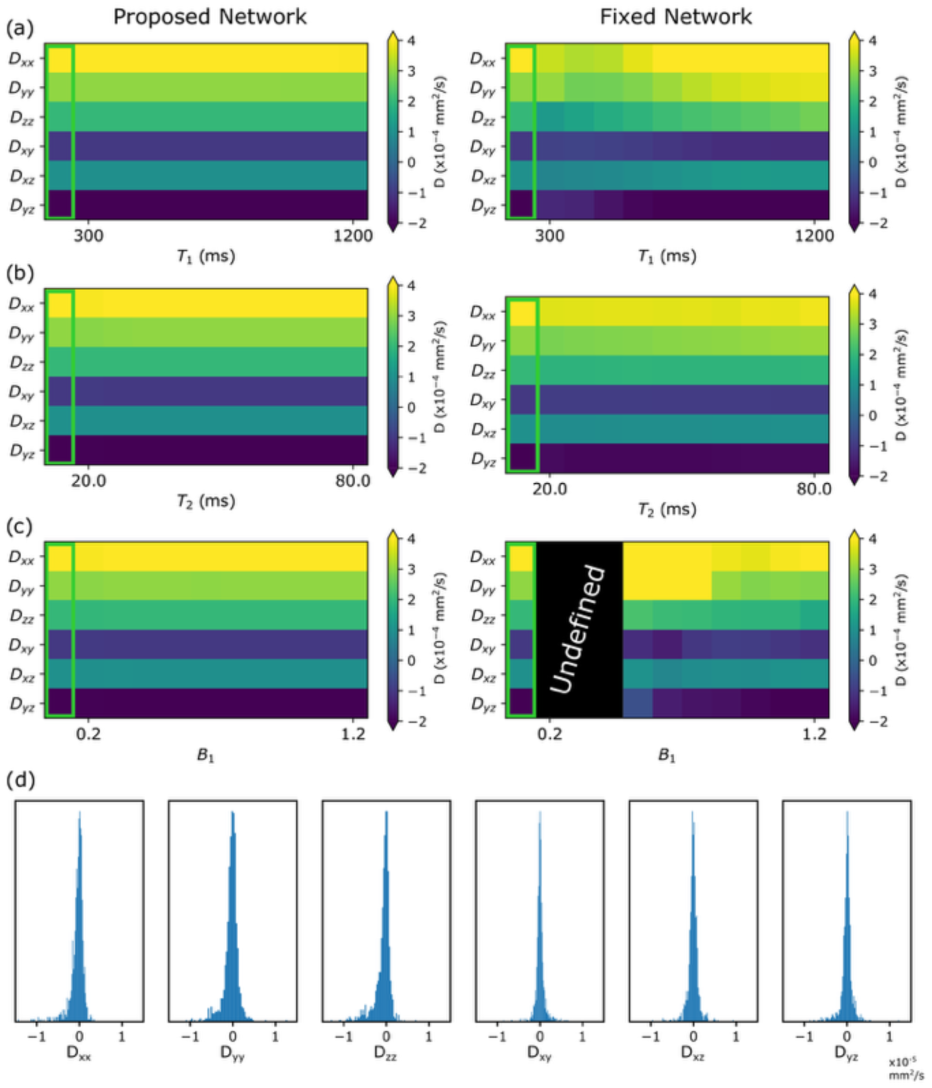
Robustness of implemented NPE network to DW-SSFP signal dependencies. (a-c) display the ground truth (green box) versus estimated Tensor coefficients as a function of (a) T_1_, (b) T_2_, and (c) B_1_, setting **D** = {4, 3, 2, −1, 1, −2} ⋅ 10^−4^ mm^2^/s with (left) and without (right) conditioning, with the latter trained on fixed relaxation times (T_1_ = 650 ms, T_2_ = 35 ms) and no transmit inhomogeneity (B_1_ =1) (see Methods). Excellent agreement is found across the dependency range incorporating conditioning (left), with median absolute difference = 5.0 ·10^-7^ mm^2^/s and interquartile range = 8.1 ·10^-7^ mm^2^/s. (d) displays the average difference (plotted as a histogram) incorporating conditioning between estimated and ground truth Tensor coefficients across 1000 simulations with arbitrary **D**, T_1_, T_2_, and B_1_ (estimated from the prior - see Methods). Excellent agreement is again found with respect to ground truth (median absolute difference = 5.7 ·10^-7^ mm^2^/s; interquartile range = 8.2 ·10^-7^ mm^2^/s). Without conditioning (a-c, right), the estimated diffusion coefficients display visible deviations from the ground truth, most notably for variations in T_1_ and B_1_, consistent with the trends in Figure 2b. 100 posterior samples were taken per simulation, with the mean of the posterior estimates used for comparison. Default values (a-c): T_1_ = 600 ms; T_2_ = 33.3 ms; B_1_ = 0.98, corresponding to simulated values closest to the mean relaxation values estimated in the acquired post-mortem experimental data and no transmit inhomogeneity (T_1_ = 650 ms; T_2_ = 35 ms; B_1_ = 1). To prevent signal scaling biasing parameter estimates when no conditioning was incorporated (right), the unconditioned signals amplitudes were normalised to then non-diffusion weighted signal amplitude corresponding to T_1_ = 650 ms; T_2_ = 35 ms; B_1_ = 1 at 24°. Undefined values (c – right) could not be estimated from the NPE network. Difference maps (a-c) provided in Supplementary Material Figure S3.

Figure 4d displays histograms of the difference between the estimated and ground truth Tensor coefficients across 1000 simulations incorporating arbitrary **D**, T_1_, T_2_, and B_1_ combinations, again finding excellent agreement (median absolute difference = 5.7 ·10^-7^ mm^2^/s; interquartile range = 8.2 ·10^-7^ mm^2^/s). Taken together, these findings demonstrate the robustness of NPE to parameter estimation within the prior range.

Figure 5a and b compares the accuracy of parameter estimation for NPE networks trained assuming non-Gaussian (a) and Gaussian (b) noise distributions, evaluated using simulated data across different SNR levels with added non-Gaussian noise and fixed **D**, with difference maps provided in Supplementary Material Figure S4. The NPE network demonstrates excellent agreement to ground truth across the SNR range (median absolute difference = 10.2 ·10^-6^ mm^2^/s; interquartile range = 8.9 ·10^-6^ mm^2^/s), with improved parameter estimation in low SNR regimes when training incorporated a non-Gaussian noise distribution. Figure 5c and d displays the residuals (difference from the ground truth) for the diagonal Tensor coefficients estimated with NPE and NLS. Here NPE residuals incorporating a non-Gaussian noise distribution display a symmetric deviation from ground truth, with the Gaussian equivalent biased towards negative values, indicative of an underestimation of Tensor coefficients. These differences reflect the potential of NPE to incorporate the underlying noise distribution when performing parameter inference, with a Gaussian noise assumption leading to biased parameter estimates arising from the impact of the noise floor (D. K. Jones & Basser, 2004). These findings are very similar to an identical evaluation using a conventional NLLS algorithm (Supplementary Material Figure S5), where parameter biases in low-SNR regimes due to the presence of non-Gaussian noise are well understood (Gudbjartsson & Patz, 1995).

**Figure 5:**
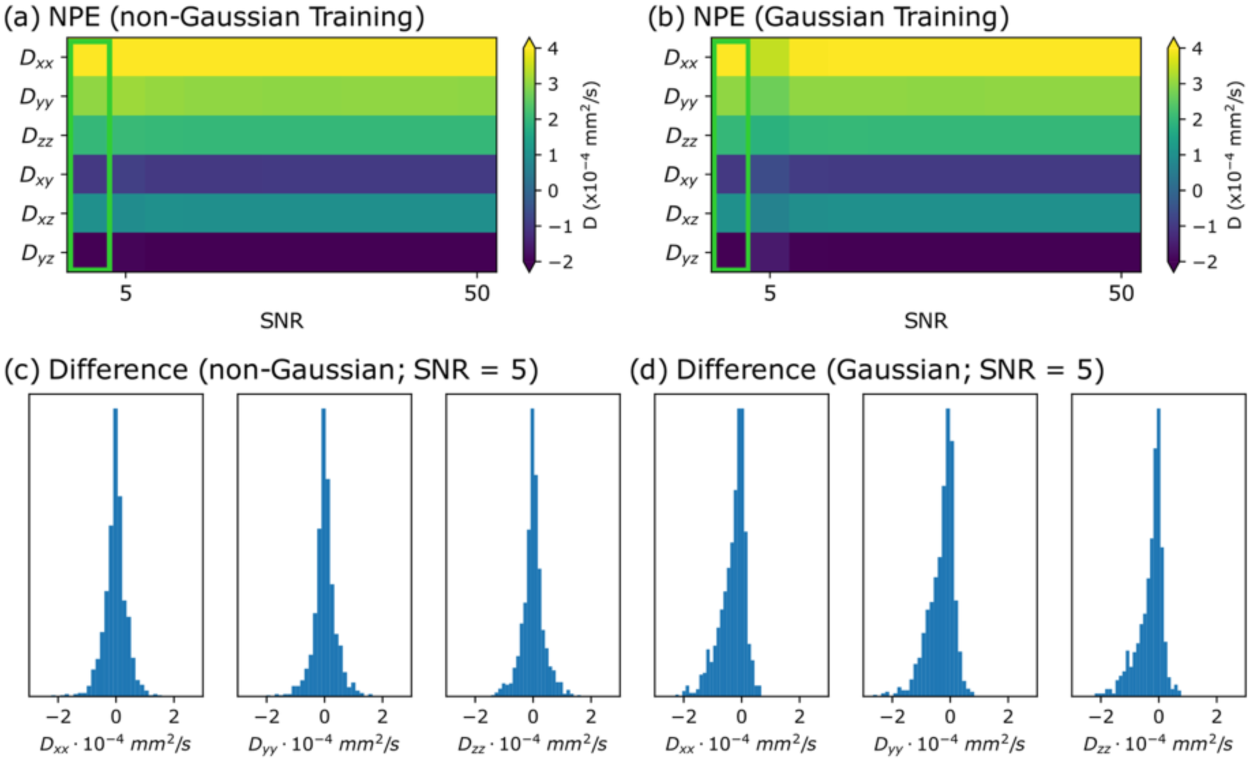
Robustness of implemented NPE network to SNR. Here I display the ground truth (green box) versus estimated Tensor coefficients for (a) an NPE network trained using simulated data incorporating a non-Gaussian noise distribution and (b) an NPE network trained using simulated data incorporating a Gaussian noise distribution. Here parameters were estimated evaluated from non-Gaussian noise-distributed DW-SSFP signals varied as a function of SNR, setting **D** = {4, 3, 2, −1, 1, −2} ⋅ 10^−4^ mm^2^/s and defining SNR with respect to the non-diffusion weighted signal. Excellent agreement is found across the SNR range (median absolute difference = 10.2 ·10^-7^ mm^2^/s; interquartile range = 8.9 ·10^-7^ mm^2^/s), reflecting the robustness of NPE to parameter estimation within the trained SNR regimes. The NPE implementation with Gaussian noise displays increased biases at low SNR values, reflecting the difference between the assumed noise model (Gaussian) and the underlying noise model of the simulated data (a noncentral chi distribution with a single degree of freedom), which are not visible using the NPE network incorporating the non-Gaussian noise distribution. These biases are visualised in (c) and (d), which display the average difference (plotted as a histogram) between estimated and ground truth diagonal Tensor coefficients across 1000 simulations with SNR = 5 and arbitrary **D**, T_1_, T_2_, and B_1_ (estimated from the prior and uniform distributions - see Methods). Whilst non-Gaussian training (c) displays symmetric residuals, Gaussian training (d) is biased towards negative values, indicative of an underestimation of **D**. Results in (a) and (b) reflect averages from parameter estimates derived from 1000 noise realisations per SNR level. Default values: T_1_ = 650 ms; T_2_ = 35 ms; and B_1_ = 1; corresponding to the mean relaxation values estimated in the acquired post-mortem experimental data and no transmit inhomogeneity. Difference maps (a and b) provided in Supplementary Material Figure S4.

Figure 6 compares the relative time performance of NLLS versus NPE as a function of the number of posterior samples. NPE can sample several thousand posterior estimates in the time required to obtain a single NLLS estimate (54.27 ms). Modelling NPE evaluation time as a linear function (Figure 6b – blue line), I estimate:

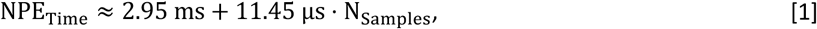

**Figure 6:**
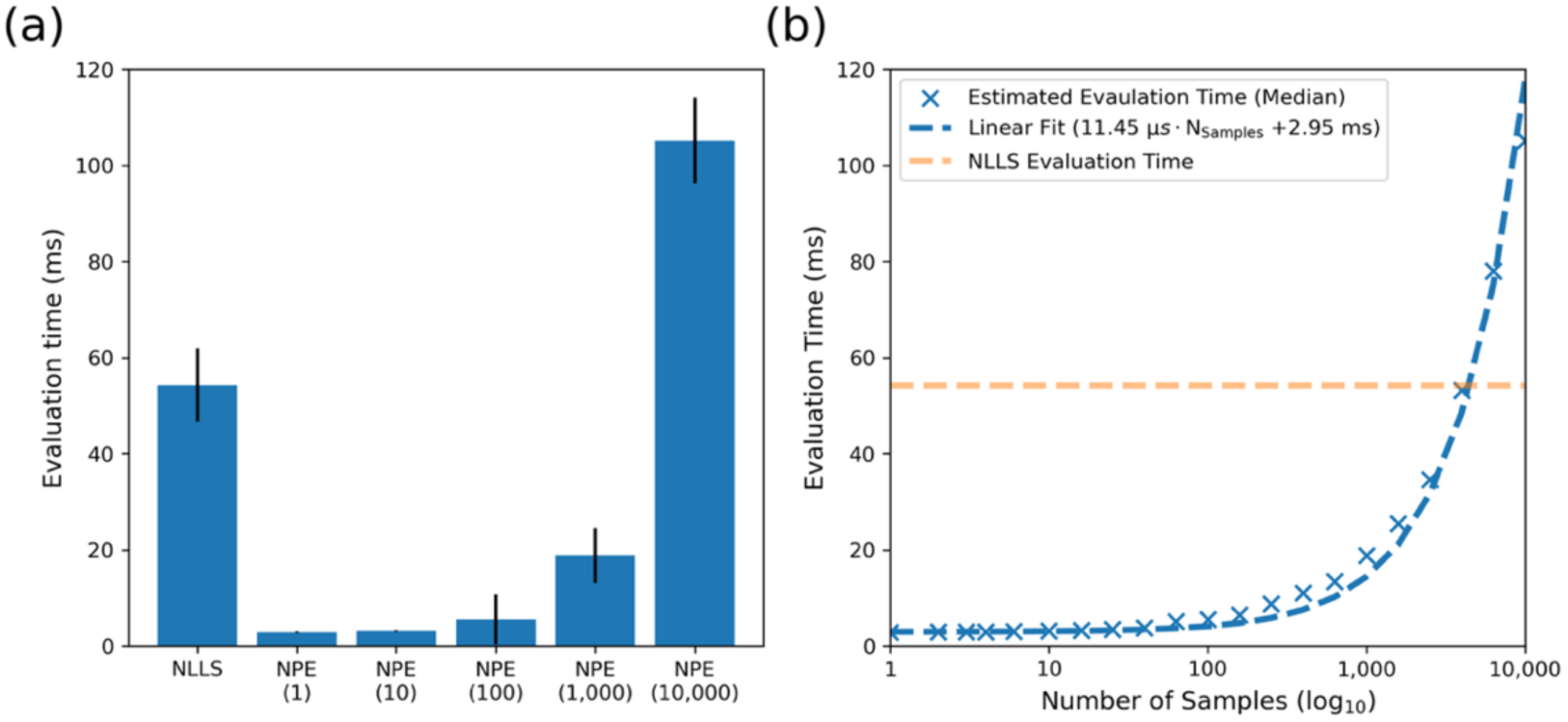
NLLS and NPE evaluation time. Simulating 1000 DW-SSFP signal profiles with arbitrary **D**, T_1_, T_2_, and B_1_ (estimated from prior and uniform distributions - see Methods) and SNR = 20 (defined relative to the non-diffusion weighted signal), (a) displays the relative evaluation time (median ± interquartile range) for NLLS and NPE as a function of the number of posterior samples (brackets along x-axis). Here we observe that we can estimate several thousand posterior samples in an equivalent NLLS evaluation time. Investigating this further (b), we can approximate evaluation time in NPE as a linear function (dashed blue line) of the number of samples and an initialisation time (offset), fitting to the median values across the sample range. Using a linear function (Eq. [1]), I estimate ≈ 4500 posterior samples in matched estimation time to NLLS. Note that the x-axis in (b) is log_10_ scale.

where N_Samples_is the number of posterior samples. Here we can interpret the offset (2.95 ms) as the initialisation time for NPE, and 11.45 µs as the approximate time required per posterior sample. Outside of the training time required for the NPE network (here ∼34 hours on a CPU), using Eq. [1] I estimate ≈ 4500 posterior samples in matched estimation time to NLLS.

For the final simulation investigation, Figure 7 displays the mean and standard deviation of the averaged Tensor coefficients as a function of the number of posterior samples and SNR (based on the DW-SSFP acquisition scheme in Table 1). Specifically, Figures 7a and b display the averaged mean and standard deviation (over tensor coefficients and simulations), with Figures 7c and d displaying the rate of change of the mean and standard deviation as a function of the number of posterior samples. The estimated mean (Figure 7a) is similar across the SNR range (consistent with Figure 5), with the standard deviation (Figure 7b) decreasing with increased SNR, consistent with our expectation of parameter uncertainty. The rate of change analysis (Figures 7c and d) indicates that a few samples (<10) are sufficient to robustly estimate the mean (defined here as <1% change), with ∼30 samples required to estimate the standard deviation. Assuming 30 samples corresponds to a ‘fully sampled’ posterior distribution, NPE provides a 16-17x acceleration versus NLLS based on the DW-SSFP acquisition scheme (Table 1) and Eq. [1].

**Figure 7:**
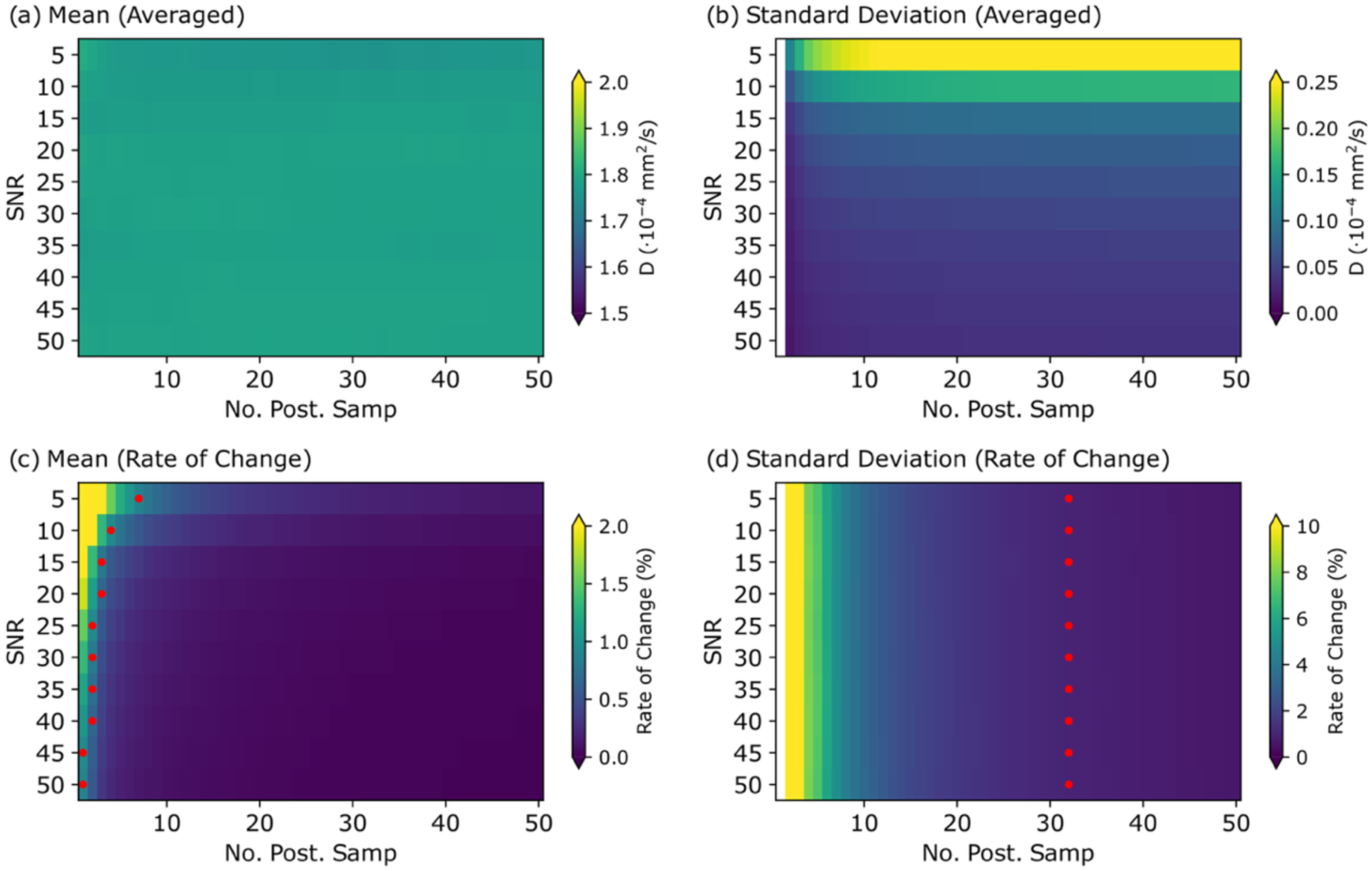
Number of samples required to characterise the posterior. Here I visualise the average (a and b) and rate of change (c and d) of the posterior mean (left column) and standard deviation (right column) as a function of the number of posterior samples (x-axis) and SNR (y-axis). The red dots in (c) and (d) correspond to the number of samples where the rate of change falls to below 1%. Figure calculated from 1000 simulations with arbitrary **D**, T_1_, T_2_, and B_1_ (estimated from the prior and uniform distributions - see Methods) using the DW-SSFP acquisition scheme described in Table 1, with estimates averaged over the Tensor coefficients and simulations. To improve visualisation and quantification, a smoothing filter (*Savitzky-Golay*) was applied to the averaged data to reduce local fluctuations. Note that the colourmaps in (a)/(b) and (c)/(d) correspond to different ranges.

### 4.2 Experimental Analysis

Figure 8 visualises the experimental Tensor estimates using NPE and NLLS. Excellent visual agreement is found for the derived Tensor outputs, with the composite chequerboard images (label in Figure 8d) displaying no visual distinctions between NPE and NLLS in the reconstructed 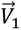 (Figure 8a), MD (Figure 8b) and fractional anisotropy (FA) (Figure 8c) maps.

**Figure 8:**
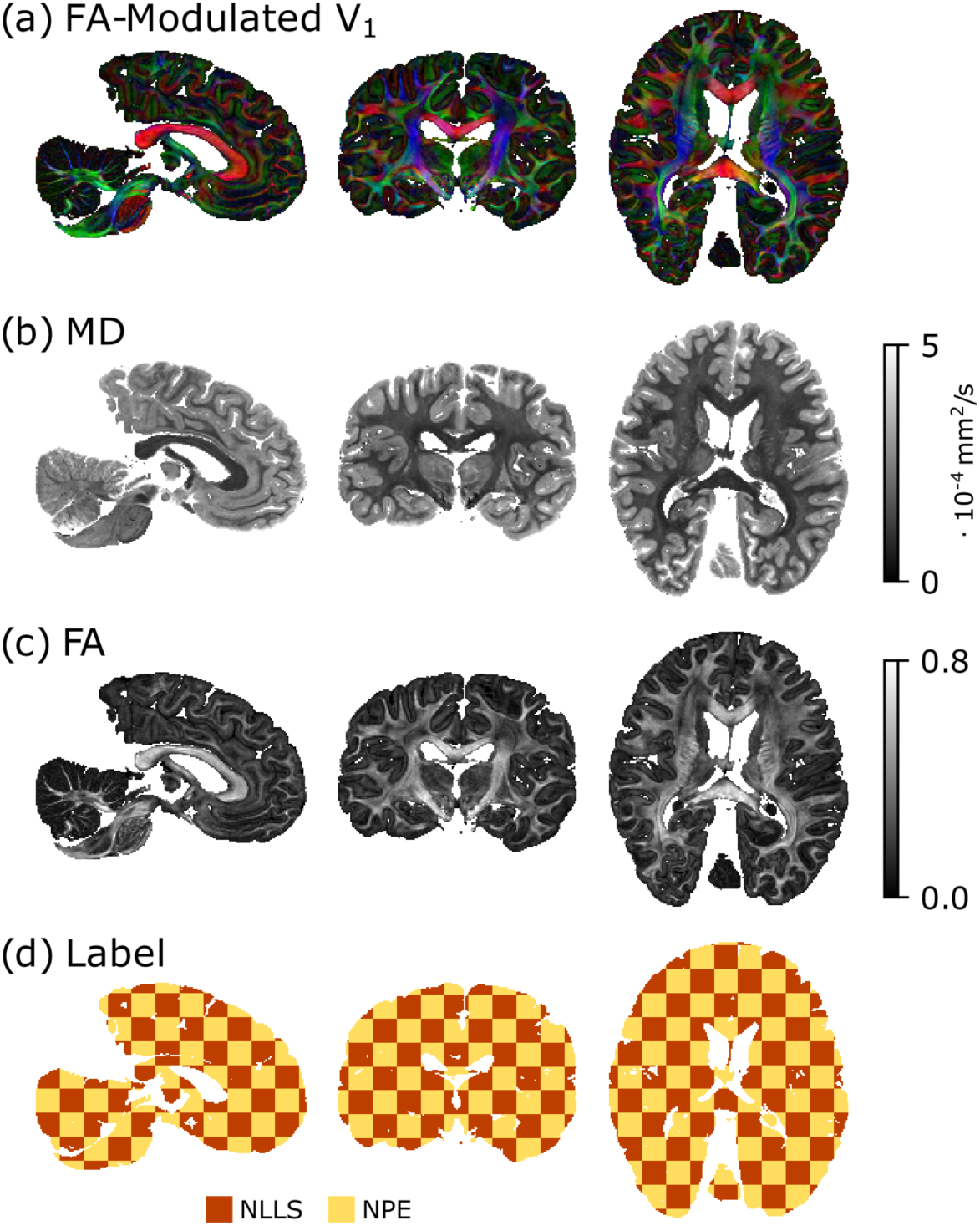
Visual comparison of experimental NPE & NLLS Tensor outputs. Composite chequerboard image (d) combining Tensor outputs from the NPE and NLLS fitting to display (a) 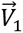 (modulated by FA), (b) MD and (c) FA maps. Excellent visual agreement is found across the brain, with no visual distinction between the NPE and NLLS outputs. Displayed NPE outputs derived from to the mean of the 100 posterior estimates per voxel.

Figure 9 quantifies the relationship between the NPE and NLLS outputs, displaying their correlation for individual Tensor elements. A strong linear relationship is found, estimating Pearson correlation coefficients r = 0.996 (D_xx_), 0.993 (D_xx_), 0.993 (D_xz_), 0.996 (D_yy_), 0.994 (D_yz_), 0.996 (D_zz_). A small bias (diagonal elements) and asymmetry (off-diagonal elements) in the correlation distribution around the unity line reflects a small underestimation of diffusion coefficients in NLLS versus NPE. This corresponds to a bias arising from non-Gaussian noise in lower SNR regions, likely arising from the assumed Gaussian noise distribution with NLLS fitting methods, or deviations between the trained noise distribution associated with the NPE network and the noise-floor corrected experimental data.

**Figure 9:**
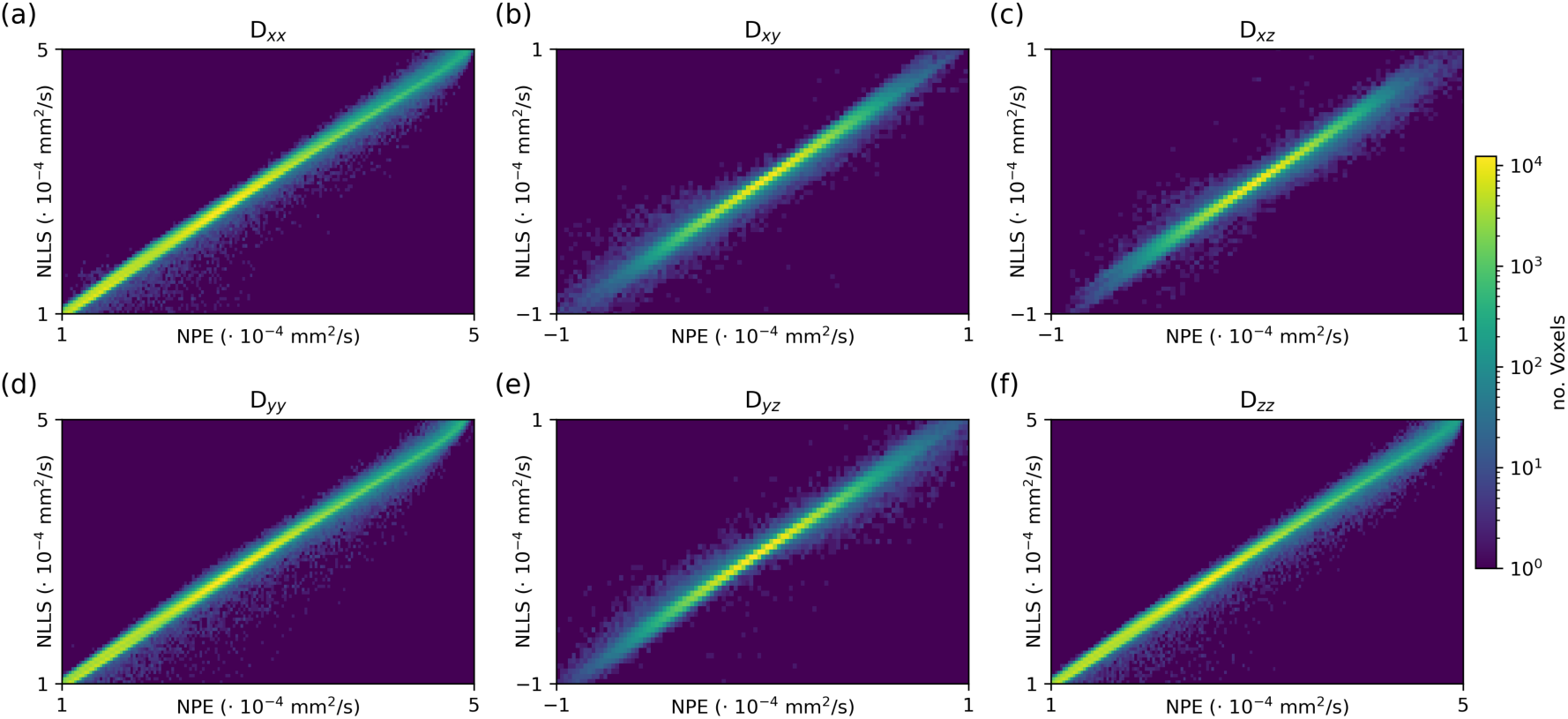
Correlation between NPE & NLLS Tensor elements. 2D histogram distributions correlating individual of NPE (x-axis) and NLLS (y-axis) Tensor elements. Excellent agreement is found across the coefficient distributions, with the histogram density defined predominantly along the diagonal. Pearson correlation coefficients for all elements >0.99. Individual histograms formed on a 200×200 grid, with the colormap reflecting a logarithmic scale (see colour bar).

A key additional advantage of NPE versus NLLS is the ability to characterise detailed posterior distributions. Figure 10 displays an example posterior distribution for a single voxel in the corpus callosum, providing detailed information characterising the precision of parameter estimates (diagonal 1D histograms) and relationship between parameters (off-diagonal 2D histograms). Tensor estimates are informative of a corpus callosum voxel, indicating a larger diffusion coefficient along a single axis most closely aligned with the fibre orientation (*D_xx_*), with smaller coefficients for orthogonal elements (*D_yy_* and *D_zz_*). The distribution was formed from 4,500 posterior samples, equivalent to the number of samples that can be achieved with a matched estimation time versus NLLS (Figure 6 and Eq. [1]).

**Figure 10:**
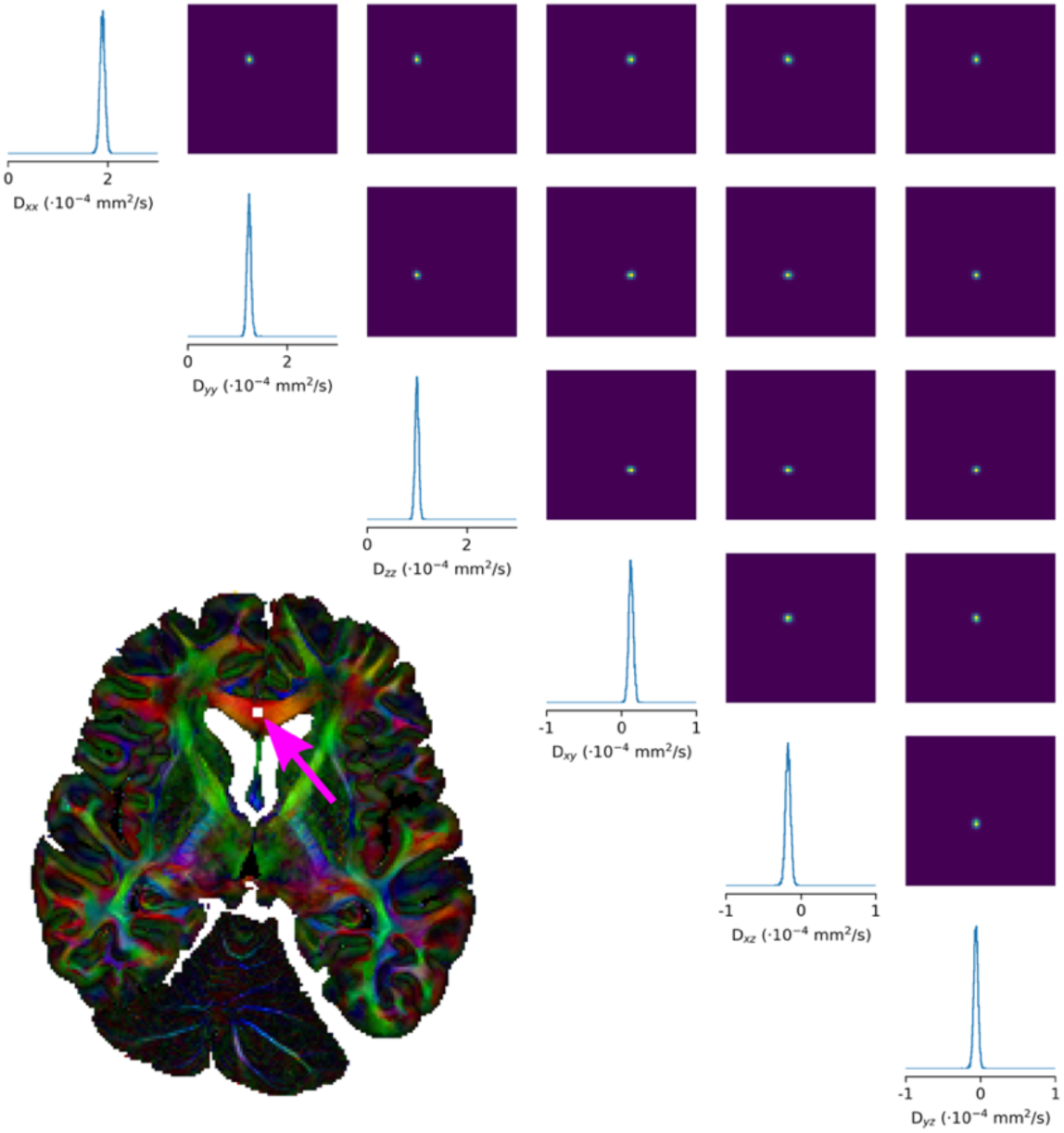
NPE posterior distribution in a single voxel. Here I visualise the NPE posterior distribution for a single experimental voxel on the corpus callosum midline (pink arrow). The diagonal (1D histograms) and off-diagonal (2D histograms) correspond to the univariate and pairwise marginal distributions. The experimental data demonstrates high confidence in the parameter estimates, with a narrow distribution of posterior samples and no degeneracy (i.e. a single peak). Posterior distribution formed from 4500 samples, equivalent to a matched estimation time versus NLLS (Eq. [1]).

Finally, Figure 11 visualises the average voxelwise signal across diffusion-weighted datasets (Figure 11a) and the corresponding standard deviation of estimated Tensor coefficients (Figure 11b). Regions of reduced signal are associated with higher standard deviations, consistent with the expectations of uncertainty quantification and findings presented in Figure 7. The majority of voxels are associated with a small standard deviation (order of 10^-5^ mm^2^/s), with higher standard deviations predominantly associated with regions close to the brain boundary along the super-inferior axis (associated with low B_1_ in this dataset) (Tendler, Foxley, Hernandez-Fernandez, et al., 2020) and deep-grey matter regions. Post-mortem deep-grey matter regions have short transverse relaxation times, leading to a very low measured DW-SSFP signal (Supplementary Material Figure S2). The use of a cartesian readout in the implemented DW-SSFP sequence leads to a small additional contribution from T_2_^’^ effects as the TE (21 ms) does not match the TR (28 ms), which leads to additional focal signal loss in iron rich deep-grey matter regions.

**Figure 11:**
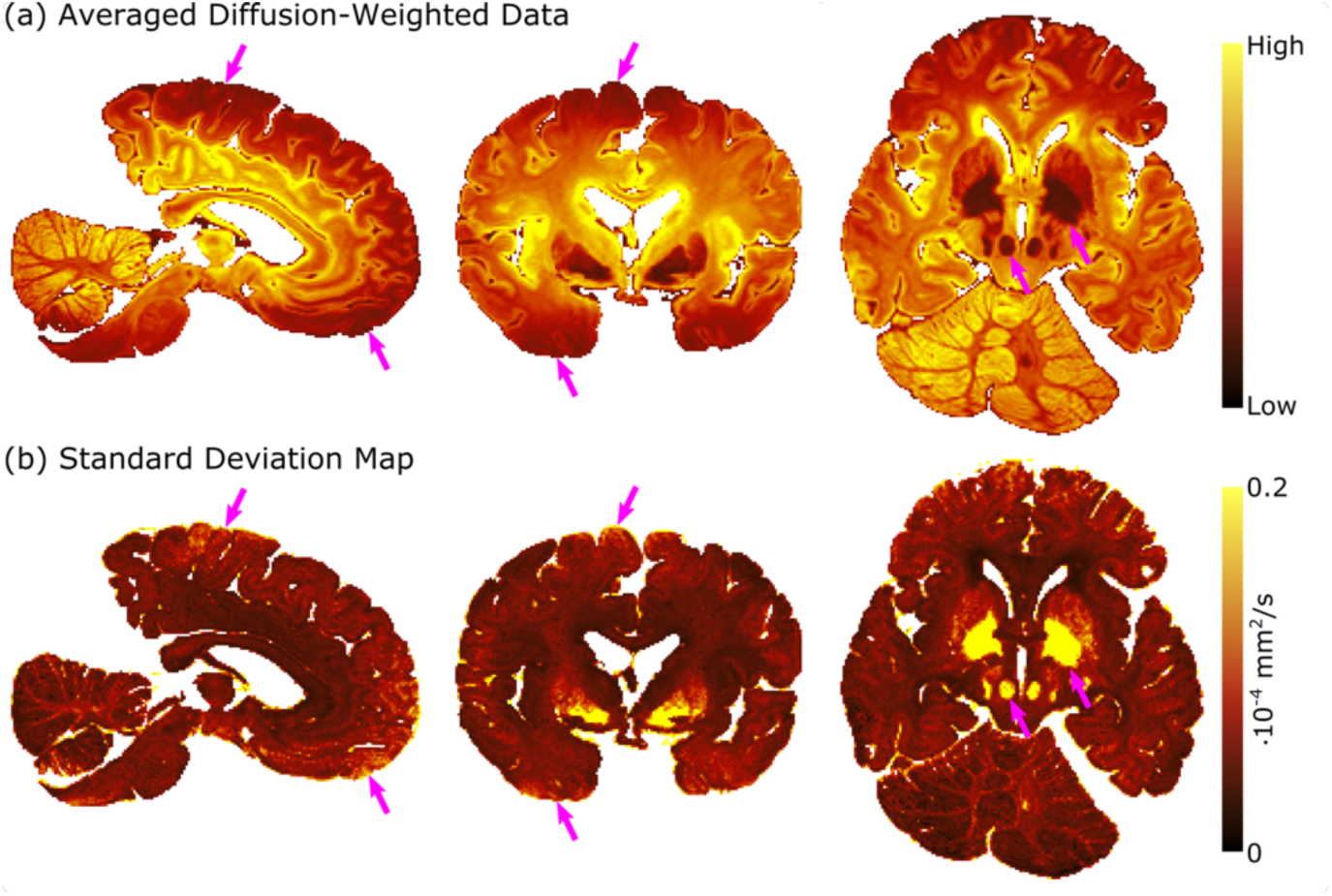
SNR and standard deviation of Tensor coefficient estimates. (a) displays the average signal amplitude across all diffusion-weighted volumes (Table 1), a proxy for SNR. (b) displays the voxelwise standard deviation of Tensor coefficients averaged over all six estimated tensor components. Voxels with a reduced signal amplitude are typically associated with a higher standard deviation, with the pink arrows highlighting example regions. The standard deviation map in (b) provides a quantitative estimate of uncertainty in parameter estimates for NPE, here defined as the average parameter standard deviation from the 100 posterior samples per voxel. Uncertainty is low (order of 10^-5^ mm^2^/s) relative to the average amplitude of diffusion coefficients per voxel (e.g. MD map in Figure 8b), with the exception of deep grey matter regions characterised by very low SNR in the acquired post-mortem data.

## 5. Discussion

### 5.1 Overview

The findings presented here demonstrate that NPE achieves fast and precise parameter inference for DW-SSFP investigations when compared to a known ground truth (synthetic data) (Figures 4 and 5) or complementary methods (NLLS) (Figures 8 and 9), with the proposed conditioning approach accounting for dependencies on T_1_, T_2_ and B_1_. NPE provides detailed parameter information, characterising full posterior distributions (Figure 10) with extremely fast inference across multidimensional parameter spaces (Figures 6 and 7), quantification of uncertainty (Figure 11), and low computational requirements.

### 5.2 SNR and noise distributions

It is well established that MRI magnitude data reconstructed from single or multiple channel coil arrays have non-Gaussian noise (typically characterised by e.g. Rician or noncentral chi distributions) (Aja-Fernández et al., 2009; Gudbjartsson & Patz, 1995). The impact of non-Gaussian noise on NPE networks was investigated theoretically using simulations, where Figure 5 demonstrates that NPE can achieve unbiased parameter estimates when trained on simulated data with added rectified noise. The impact of non-Gaussian noise on NLLS fitting methods is well understood, and for the experimental analysis non-Gaussian noise effects were attenuated via a noise-floor correction (Gudbjartsson & Patz, 1995) to facilitate comparisons between NPE and NLLS. Whilst this may lead to noise distribution biases, the experimental acquisition (described in in (Tendler, Foxley, Hernandez-Fernandez, et al., 2020)) was designed to ensure all experimental voxels are associated with some high-SNR data, with excellent experimental parameter agreement between NPE and NLLs with different assumed noise distributions (Figures 8 and 9) indicating that any biases are small.

When considering different noise distributions (e.g. theoretically calculated or estimated from experimental data), these could be similarly incorporated into the forward simulation model for NPE training. Alternatively, NPE networks could be conditioned on noise floor estimates via *P*(θ | S, *T*_1_, *T*_2_, *B*_1_, S_nf_), incorporated into the forward simulation model via *S*_measured_ = (*S*^2^ + S^2^)^1/2^ (Gudbjartsson & Patz, 1995). Experimental noise floor estimates (i.e. calculated from the background signal) could then be passed to the network during parameter inference to reduce parameter biases. Datasets could be alternatively denoised (e.g. using PCA based methods) to reduce parameter biases as part of a standard pre-processing pipeline prior to parameter estimation.

### 5.3 Deviations from Tensor representation

Whilst the synthetic training data precisely characterises the DW-SSFP signal under a Tensor representation, it is well established that experimental diffusion MRI data can be characterised by a more sophisticated orientation function. One example of this is in crossing fibre regions, characterised by a low FA when fitting a single Tensor to experimental data. This raises a question of whether NPE will fail when performing parameter estimation in tissue regions that have signal representations that deviate substantially from the training data, or whether the detailed posterior distributions will demonstrate sensitivity to multiple fibre populations.

To investigate whether such effects were present in the current dataset, NPE Tensor estimates were investigated in the centrum semiovale (Figure 12), a white matter crossing fibre region. In both NPE and NLLS this region was characterised by a low FA (Figure 12a and b), with no evidence of NPE distinguishing individual fibre populations in the posterior distribution (Figure 12c). Specifically, NPE consistently identified a single peak, with estimated diffusion coefficients characteristic of isotropic diffusion (consistent with NLLS). Whilst previous work has demonstrated multimodal posterior distributions under a Ball and Sticks signal representation (Manzano-Patron et al., 2025), one interpretation is that the underlying Tensor fit to experimental data is not truly degenerate (i.e. characterising diffusivity as more isotropic is a better representation of the experimental data versus accurately characterising one of the Tensors), which can be effectively captured by NPE. Notably, we do expect orientation-distributions in post-mortem DW-SSFP data to be sensitive multiple fibre populations (Tendler, Foxley, Hernandez-Fernandez, et al., 2020, Tendler, Warrington, et al., 2025).

**Figure 12:**
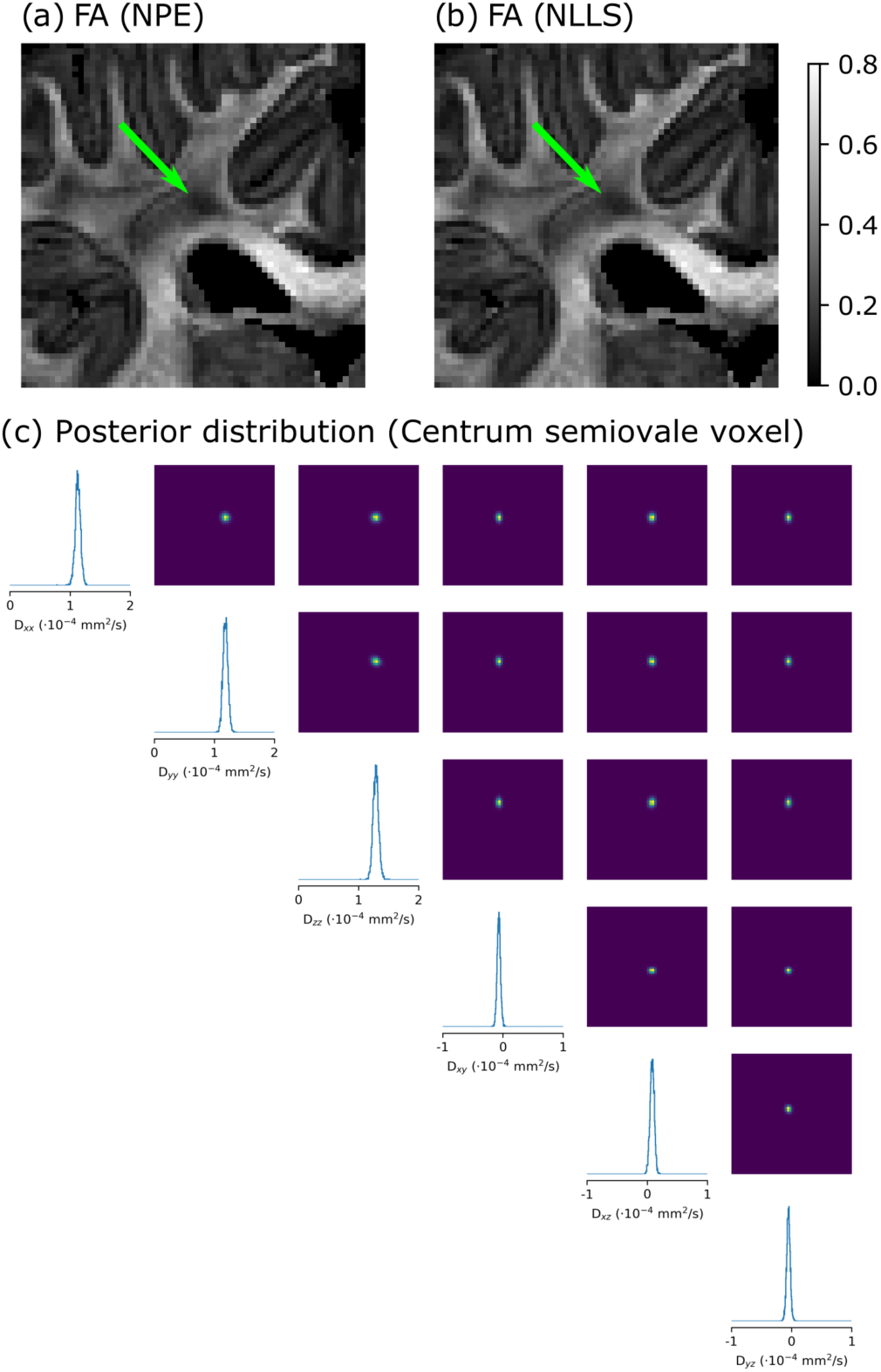
Comparisons of FA estimates in the centrum semiovale. Here I visualise NPE (a) and NLLS (b) estimates of FA in the centrum semiovale of the post-mortem brain, a white matter crossing fibre region. Both NPE and NLLS characterised the region with low FA (green arrow). Posterior distributions estimated from NPE, displayed in (c) for a single centrum semiovale voxel with low FA, were characteristic of isotropic diffusion, with similar diffusion coefficients across the diagonal elements (*D_xx_*, *D_yy_*, *D_zz_*) and low diffusivity in the off-diagonal elements (*D_xy_*, *D_xz_*, *D_yz_*). (c) formed from 4,500 posterior samples.

Characterisation of non-Gaussian/multimodal posterior distributions estimated with NPE is an area of active investigation (Jallais & Palombo, 2024). To further investigate deviations from Gaussian-distributed posteriors in the post-mortem data investigated here, Figure 13 compares FA estimates calculated from the mean and maximum-a-posteriori (MAP) of the posterior distribution. No systematic difference were identified in the parameter estimates, indicative that the posterior distributions are Gaussian, with the exception of deep grey matter regions characterised by extremely low SNR in this dataset (Figure 11).

**Figure 13:**
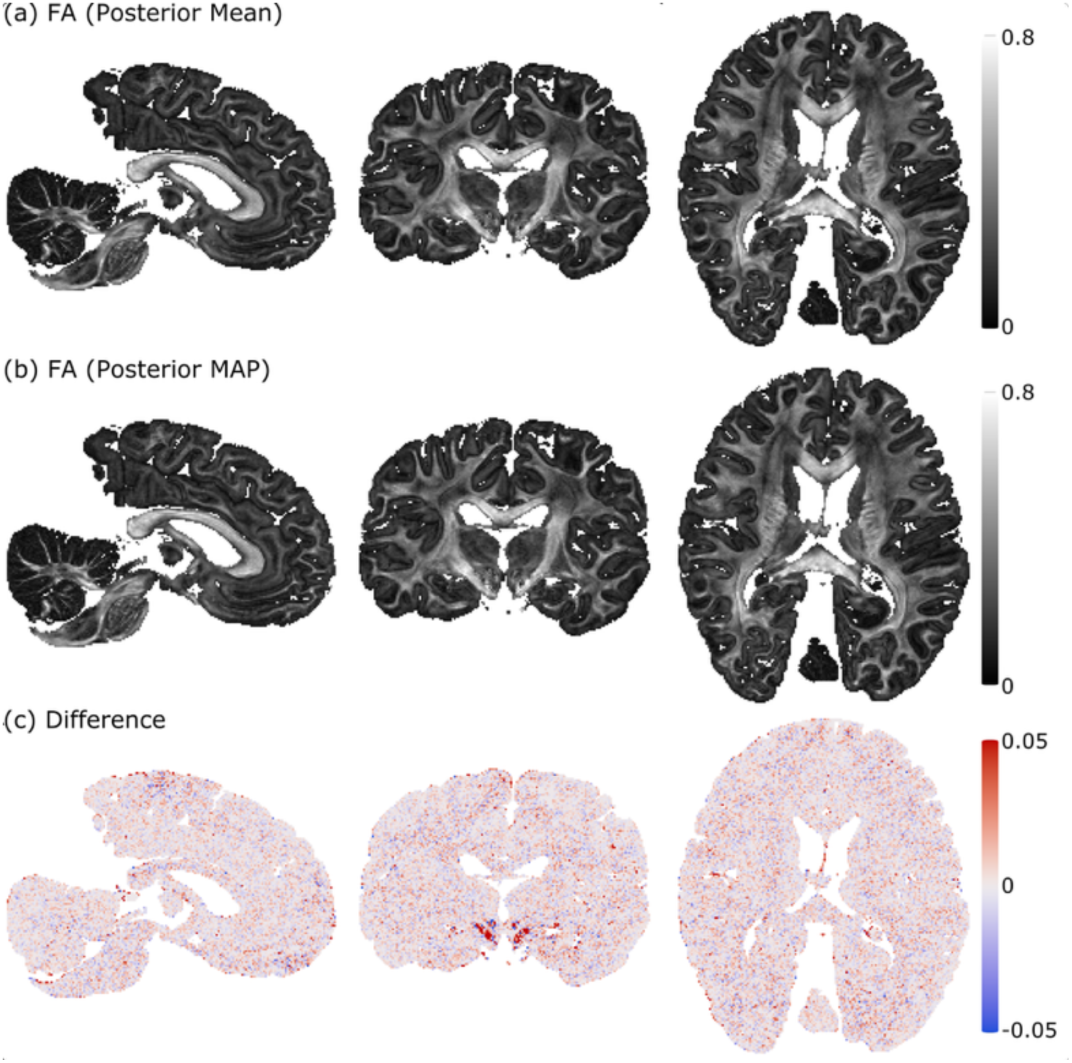
Mean and MAP FA estimates. Here I visualise the posterior mean (a) and MAP (b) estimates of FA across the whole brain. The two parameter maps display excellent agreement, with no systematic differences across anatomy (c), except for deep grey matter regions characterised by a very low SNR in this dataset (Figure 11). MAP estimates calculated by evaluating a Gaussian kernel density estimator (KDE) over the posterior-distribution, with the sample corresponding to the highest probability density estimate selected as the MAP value.

### 5.4 Limitations

One limitation of NPE for diffusion MRI investigations is that networks typically incorporate a specific forward model, defined by a fixed number of parameters or signal estimates. Whilst changes in the diffusion MRI acquisition scheme (e.g. gradient orientations, number of directions, number of shells) can be trivially incorporated with conventional iterative fitting methods (e.g. NLLS, MCMC), in NPE these changes can require a different network architecture. At one extreme, this requires full retraining of the network to account for sequence parameter changes. Such challenges can arise even in the circumstance of missing data (e.g. arising from corrupted diffusion MR volumes).

When considering missing data, imputation methods can be introduced to approximate the missing information. This can be performed by the user (e.g. interpolation to approximate the signal arising from a missing diffusion MRI volume), or leveraging advances in SBI to jointly optimise a data imputation and NPE network (Verma et al., 2025). When considering changes in b-vectors (i.e. orientation or number), dimensionality reduction techniques can also be utilised to condense the acquired data into a subset of orientation-invariant data ‘features’ prior to NPE. These can be similarly defined by the user (e.g. spherical harmonics) (Descoteaux et al., 2007; Manzano-Patron et al., 2025) or using a joint dimensionality reduction and NPE network architectures (Radev et al., 2023; Jallais & Palombo, 2024). Addressing such challenges is an area of active investigation in NPE.

The proposed conditioning method introduced in this manuscript may similarly facilitate the generalisability of NPE networks for diffusion MRI investigations. Specifically, NPE networks could be trained over a range of signals arising from different sequence parameters combinations, with the sequence parameters provided to the NPE network for conditioning. For example, considering a DW-SE investigation, an NPE network could be conditioned on the b-value via *P*(θ | S, *b*), circumventing the requirement of explicitly training a different network per b-value. Such conditioning could be combined with missing data and dimensionality reduction techniques to improve the generalisability of NPE networks across acquisition schemes and datasets.

Differences in relaxation times can vary substantially in post-mortem tissue both within (e.g. due to tissue type) and across (e.g. due to choice of fixative) samples, motivating the acquisition of voxelwise maps for precise quantification. Spatial B_1_ distributions have known variations across acquisitions due to the choice of field strength, RF hardware and sample size. However, a limitation of DW-SSFP for diffusion coefficient estimation is the assumption that estimated experimental relaxation and B_1_ values correspond to a known ground truth, whereas differences in these estimates could bias the calculated diffusion coefficients. Such differences could arise from multiple sources, including partial volume effects associated with changes in spatial resolution across modalities (as present in this manuscript), the choice of acquisition sequence (e.g. variable flip angle or inversion recovery/spin echo methods) or sequence timings, which can lead to different parameter values. Figure 2b provides an insight into the extent of these biases for a given set of DW-SSFP parameters. For example, we can see for the DW-SSFP sequence parameters investigated in Figure 2b that we are relatively insensitive to biases in T_2_, and more sensitive to biases in B_1_ and T_1_, although this relationship is expected to vary depending on the choice of acquisition parameters. Such information could help inform where to spend most time acquiring dependency maps, or quantification of additional sources of error on diffusion coefficient estimation. The acquired dependency maps could further help inform the distribution of T_1_/T_2_/B_1_ values passed to the NPE network during training (e.g. representative of the experimental data rather than uniform distributions used in this manuscript) to improve parameter accuracy or reduce training requirements, with the limitation that the trained NPE network may be less generalisable to other datasets. Where relaxation and transmit-inhomogeneity maps have been incorporated into a DW-SSFP investigation, I recommend the reporting of the methods used in the manuscript to facilitate comparability.

### 5.5 Future directions

Whilst the work presented here aims to demonstrate the feasibility of NPE for DW-SSFP investigations by utilising a signal representation that can be evaluated using complementary methods (e.g. Tensor and NLLS), the implemented routine could be extended to incorporate more sophisticated microstructural models, including Monte Carlo simulations and alternative DW-SSFP signal representations that incorporate time-dependence (Tendler, 2025). Whilst a Tensor representation has a relatively simple posterior distribution (typically characterised by a single symmetric peak in a multidimensional space), previous work has demonstrated that NPE can incorporate sophisticated posterior distributions, with demonstrated applications for the DW-SE sequence (Eggl & De Santis, 2024; Jallais & Palombo, 2024; Manzano-Patron et al., 2025). The work presented here provides a natural extension for DW-SSP investigations, or alternative steady-state diffusion MRI sequences, including fingerprinting methods.

## 6. Conclusion

NPE demonstrates fast and precise parameter inference for DW-SSFP investigations when compared to complementary fitting methods or synthetic data with a known ground truth. The proposed method provides an intuitive approach to incorporate conditional dependencies with NPE, successfully accounting for dependencies of the DW-SSFP signal on relaxation times and transmit inhomogeneity. Presented findings provide a framework for integrating advanced microstructural models with steady-state diffusion MRI sequences, generating detailed posterior distributions for the characterisation of parameter estimates, uncertainty and degeneracies.

## Data and Code Availability

Software for the NPE implementation, including scripts to replicate many of the findings presented in this manuscript, are available at github.com/BenjaminTendler/SBI_DWSSFP. The original processed post-mortem data associated with this project can be accessed via the Digital Brain Bank (open.win.ox.ac.uk/DigitalBrainBank) (Tendler et al., 2022) (*Human ALS MRI-Histology dataset*).

## Author Contributions

BCT was responsible for all aspects of the investigation.

## Acknowledgements

BCT would like to thank Jose Pedro Manzano Patron for helpful discussions during the development of this project and manuscript feedback, and Saad Jbabdi for manuscript feedback. Human tissue was provided by the Oxford Brain Bank, funded by the Medical Research Council, Brains for Dementia Research, and the NIHR Oxford Biomedical Research Centre.

## Appendix 1

Based on the proposed analytical model in (Freed et al., 2001), the DW-SSFP sequence incorporating a diffusion gradient of duration *δ* is modelled as:

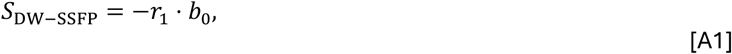

where:

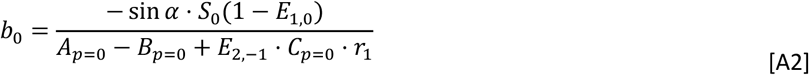

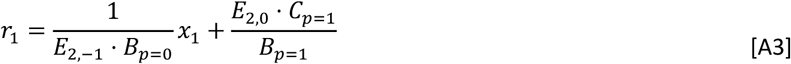

defining:

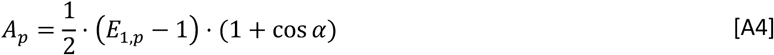

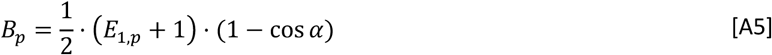

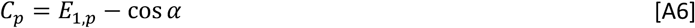

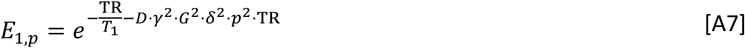

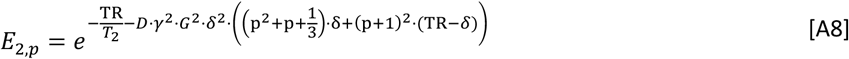

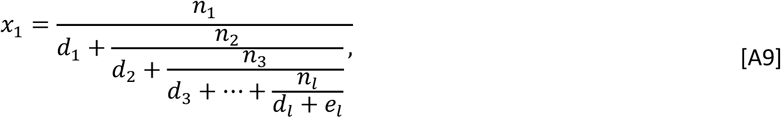

with:

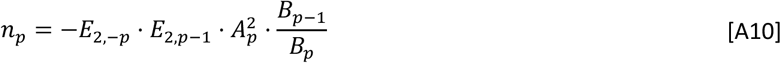

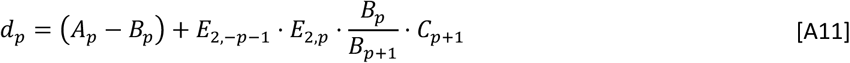

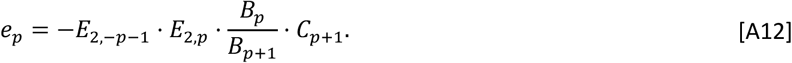

Transmit (B_1_) inhomogeneity can be incorporated as a modulation of the nominal flip angle *α*.

## Appendix 2

The DW-SSFP analytical model with integrated diffusion tensor is equivalent to Appendix 1, setting:

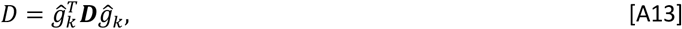

where *ĝ_k_* is the gradient orientation and ***D*** is the diffusion tensor.

