## Supplementary Material for "Integration of steady-state diffusion MRI with Neural Posterior Estimation (NPE) for post-mortem investigations"

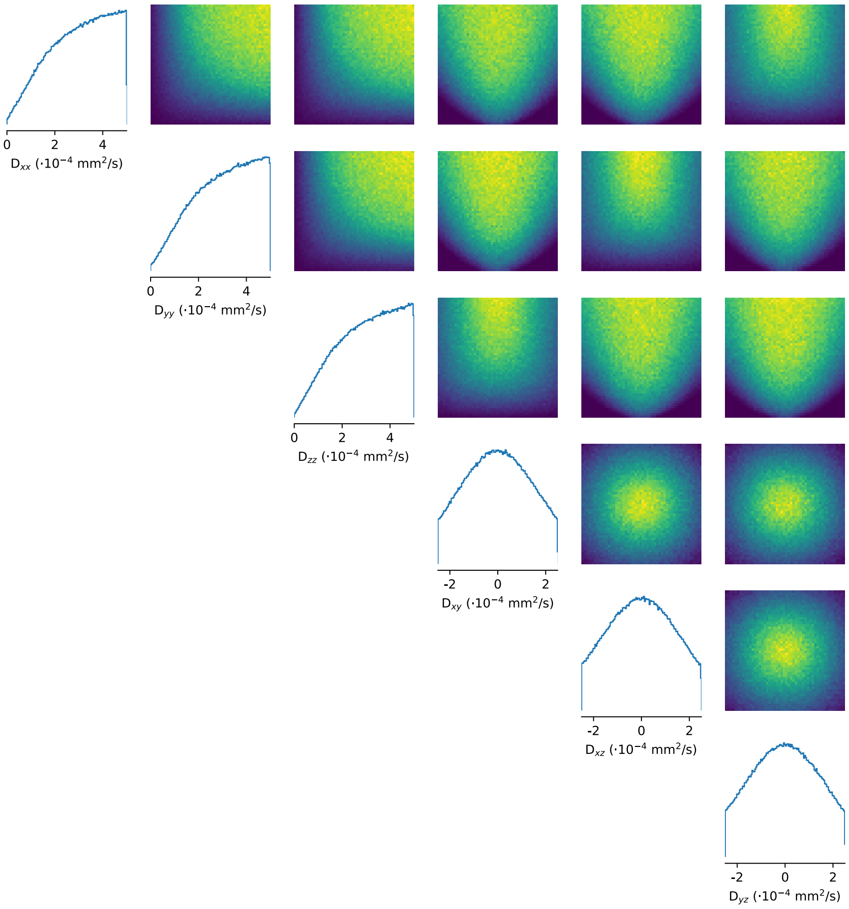


Figure S1: **Prior distribution following incorporation of the *RestrictionEstimator* function**. Prior distribution output by the trained a classifier network to facilitate the synthesis of sampled parameter combinations corresponding to positive semi-definite tensors. The diagonal (1D histograms) and off-diagonal (2D histograms) correspond to the univariate and pairwise marginal distributions.

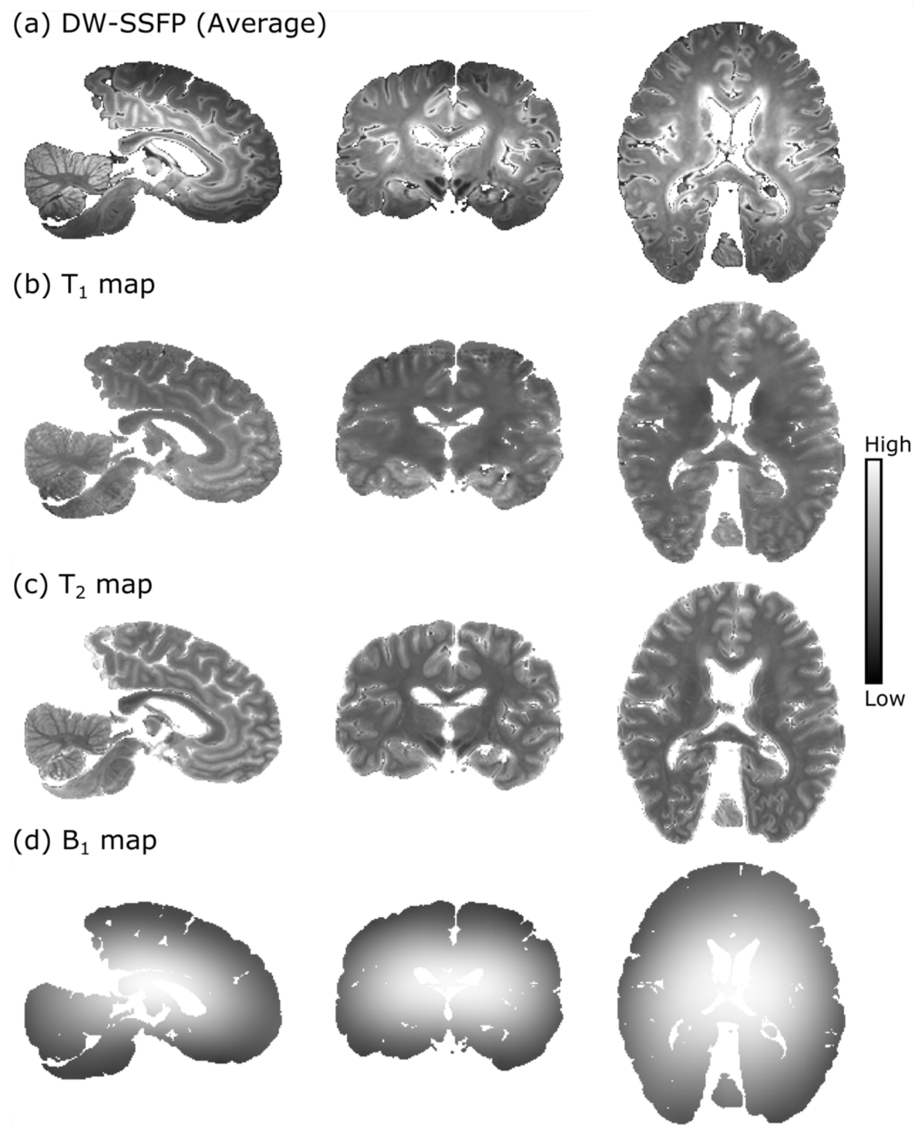


**Figure S2: DW-SSFP data and dependency maps**. Here we display the average of the DW-SSFP data (a) across diffusion and non-diffusion weighted volumes, alongside dependency maps (b-d). Data were coregistered using a six degrees of freedom coregistration only, where the low-distortion DW-SSFP data displaying excellent geometric consistency. Here the $T_1$ map is scaled between 0 and 1500 ms, the $T_2$ map is scaled between 0 and 75 ms, and the $B_1$ map is scaled between 0 and 1.2 (normalised units).

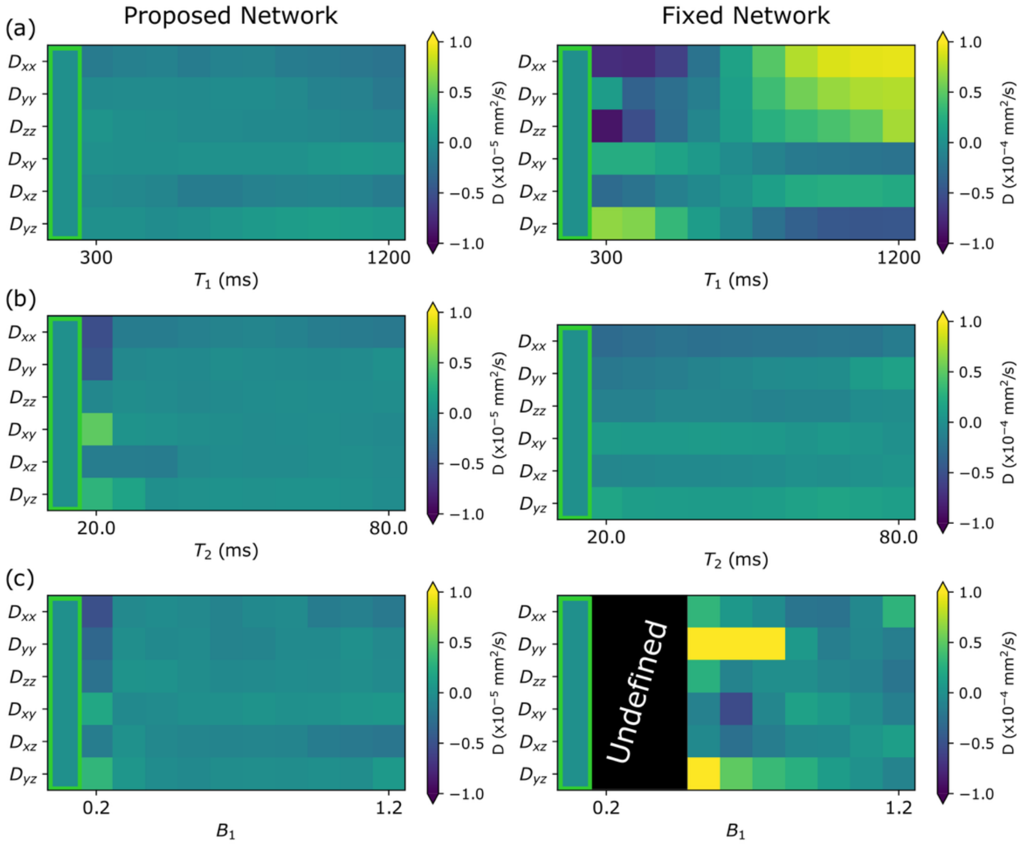


**Figure S3: Differences from ground truth (proposed and fixed network).** Equivalent to the format of Figure 4a-c (Main Text), here I display the deviation of parameter visualisations from ground truth. Note the differences in scaling of the colour bars ($10^{-4}$ $\mathrm{mm}^2$/s for the fixed network in comparison to $10^{-5}$ $\mathrm{mm}^2$/s for the proposed network).

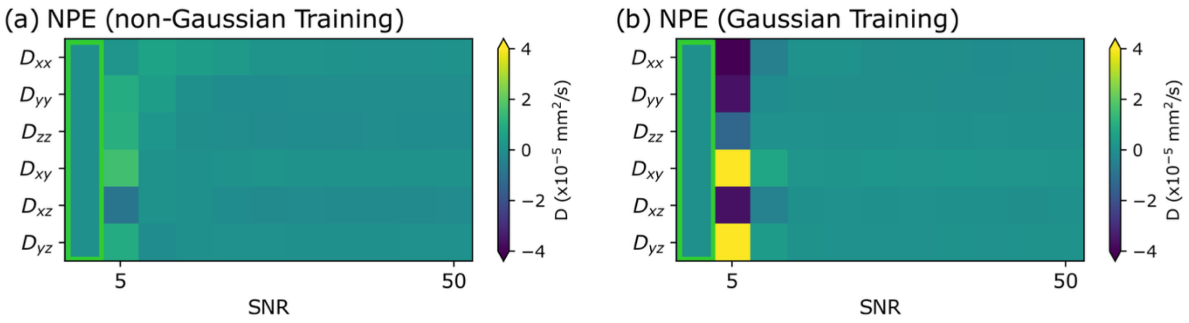


**Figure S4: Differences from ground truth (non-Gaussian and Gaussian network).** Equivalent to the format of Figure 5 (Main Text), here I display the deviation of parameter visualisations from ground truth.

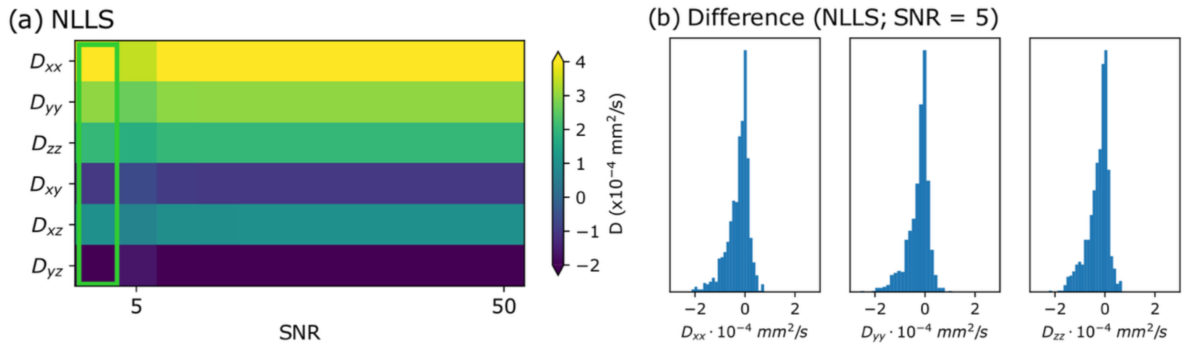


**Figure S5: Robustness of NLLS network to SNR.** Here I display the ground truth (green box) versus estimated Tensor coefficients for (a) an NLLS network trained using simulated data incorporating a non-Gaussian noise distribution. Here parameters were estimated evaluated from non-Gaussian noise-distributed DW-SSFP signals varied as a function of SNR, setting $\mathbf{D} = \{4, 3, 2, -1, 1, -2\} \cdot 10^{-4}\ \mathrm{mm/s}$ and defining SNR with respect to the non-diffusion weighted signal. The NLLS implementation displays increased biases at low SNR values, reflecting the difference between the assumed noise model (Gaussian) and the underlying noise model of the simulated data. These biases are visualised in (b), which display the average difference (plotted as a histogram) between estimated and ground truth diagonal Tensor coefficients across 1000 simulations with SNR = 5 and arbitrary $\mathbf{D}$, $T_1$, $T_2$, and $B_1$ (estimated from the prior and uniform distributions - see Methods). Results in (a) reflect averages from parameter estimates derived from 1000 noise realisations per SNR level. Default values: $T_1$ = 650 ms; $T_2$ = 35 ms; and $B_1$ = 1; corresponding to the mean relaxation values estimated in the acquired post-mortem experimental data and no transmit inhomogeneity.
